# Sunflower sea star (*Pycnopodia helianthoides*) environmental DNA (eDNA) shedding and decay: Influence of sunlight and comparison to other marine organisms

**DOI:** 10.64898/2026.10.15.759472

**Authors:** Meghan M. Shea, Laura Kubiatko, Andrew Kim, Ashley Kidd, Zachary Gold, Alexandria B. Boehm

## Abstract

Conservation practitioners increasingly turn to environmental DNA (eDNA) for biomonitoring, yet interpreting DNA-based data requires attention to organismal shedding and decay dynamics. Here, we characterize shedding rates and decay rate constants of eDNA from *Pycnopodia helianthoides,* a critically endangered and ecologically important sea star in the Northeast Pacific Ocean. Shedding rates were quantified via a mesocosm experiment utilizing twelve stars of varying sizes, and decay rate constants were calculated for naturally shed *P. helianthoides* eDNA under natural sunlit and dark conditions for 96 hours. We additionally completed a systematic search and review of marine organism shedding rates and decay rate constants. *P. helianthoides* decay rate constants (k) were not different under sunlit and dark conditions (average k=0.0257 hr^-1^) and among the lowest third reported for marine organisms. Similarly, biomass-normalized shedding rates (between 896 and 19,800 copies/hr/g) and shedding rates per individual (between 22,000 and 548,000 copies/hr/individual) were among the lowest recorded across marine organisms. Shedding rates were influenced, in part, by organism stress levels. This work aids interpretation of measured eDNA concentrations in marine environments and serves as a roadmap for integrating shedding and decay studies into eDNA assay design.

## Introduction

Kelp forests are ecologically, economically, and culturally vital ecosystems ^1^; however, due to human impacts ranging from coastal pollution^2–5^ to climate stressors^6–8^, kelp forests are declining at an estimated rate of 2% per year globally^8^. The decline has been especially prominent along the Pacific Coast of North America over the past decade, where widespread loss of kelp forests has been linked to a trophic cascade beginning with an outbreak of sea star wasting disease in 2013^9^. Sea star wasting disease, a causative agent of which has recently been identified as *Vibrio pectenicida* strain FHCF-3^10^, killed billions^11^ of sea stars across 20 species^12^ representing one of the largest marine mass mortalities to date^13^. While some sea star species have begun to recover, *Pyconopodia helianthoides* (the sunflower sea star) has not. *P. helianthoides* lost more than 50% of its range and more than 90% of its historic population size, leading to its recent International Union for Conservation of Nature (IUCN) listing as critically endangered^11^.

The decline of *P. helianthoides* is likely exacerbating kelp forest degradation. *P. helianthoides* are voracious consumers of purple sea urchins^14^; with the loss of this top benthic predator, urchin populations have exploded, and their increased grazing on kelp has transformed healthy ecosystems into urchin barrens^9^. *P. helianthoides* influences kelp predation indirectly, too, by releasing a chemical cue that creates a “landscape of fear” that suppresses urchin grazing^15^.

Because of these consumptive and non-consumptive impacts, there is significant interest in locating existing *P. helianthoides* refugia and also enabling their reintroduction across their historic range^9^.

However, these conservation goals require new monitoring approaches. Despite an expansion of visual monitoring efforts, it remains impractical to adequately survey the full range of *P. helianthoides*: from Baja Sur California, Mexico to Alaska, USA and across a depth range of over 400 meters^16^. Targeted environmental DNA (eDNA) approaches may provide a viable alternative for expanding *P. helianthoides* monitoring capacity. eDNA refers to the genetic material found in a sample of soil, sediment, water, air, or another environmental matrix^17^; this DNA, which can be intracellular or extracellular, results from sources including shed epithelial cells, gametes, metabolic waste, and tissue from decomposing organisms^18^. While some studies use sequencing approaches to detect a wide range of organismal eDNA in a sample^19^, targeted eDNA approaches use quantitative PCR (qPCR), digital PCR, or CRISPR-based methods to identify and quantify organism-specific DNA^20^. Targeted eDNA approaches have a variety of potential benefits for marine organism biomonitoring; a recent systematic review of studies comparing conventional and eDNA approaches found that eDNA surveys are more likely to yield higher detection probabilities than conventional sampling, while simultaneously lowering cost and sampling effort^21^. In response to these benefits, an eDNA assay targeting the *nad5* gene region of *P. helianthoides* was recently developed and tested in laboratory, mesocosm, and field settings^22^.

Utilizing the *P. helianthoides* assay for biomonitoring requires additional context and validation; how can we infer the density and location of *P. helianthoides* from just the concentration and/or presence of their DNA in the environment? Answering this question requires attention to shedding (how much DNA is produced per unit time), transport (how DNA is advected and dispersed over time), and fate (how quickly DNA decays over time)^23,24^ (Figure 1). Significant research over the past decade has characterized eDNA shedding rates and decay rate constants for a variety of marine organisms and environmental conditions^24–26^. Shedding rates and decay rate constants can vary by multiple orders of magnitude and are influenced by a variety of biotic and abiotic factors, including organism morphology^25,27^, water temperature^25^, and DNA fragment length^28^. Additionally, eDNA decay dynamics have been hypothesized to be influenced by photochemistry^29^, although the few studies focused on the role of sunlight on eDNA decay in marine environments have not captured this effect^30,31^. The complexity of eDNA shedding and decay necessitates species-specific studies in cases such as this one, where the ultimate goal is to operationalize a large-scale eDNA monitoring program for a rare organism.

**Figure 1:**
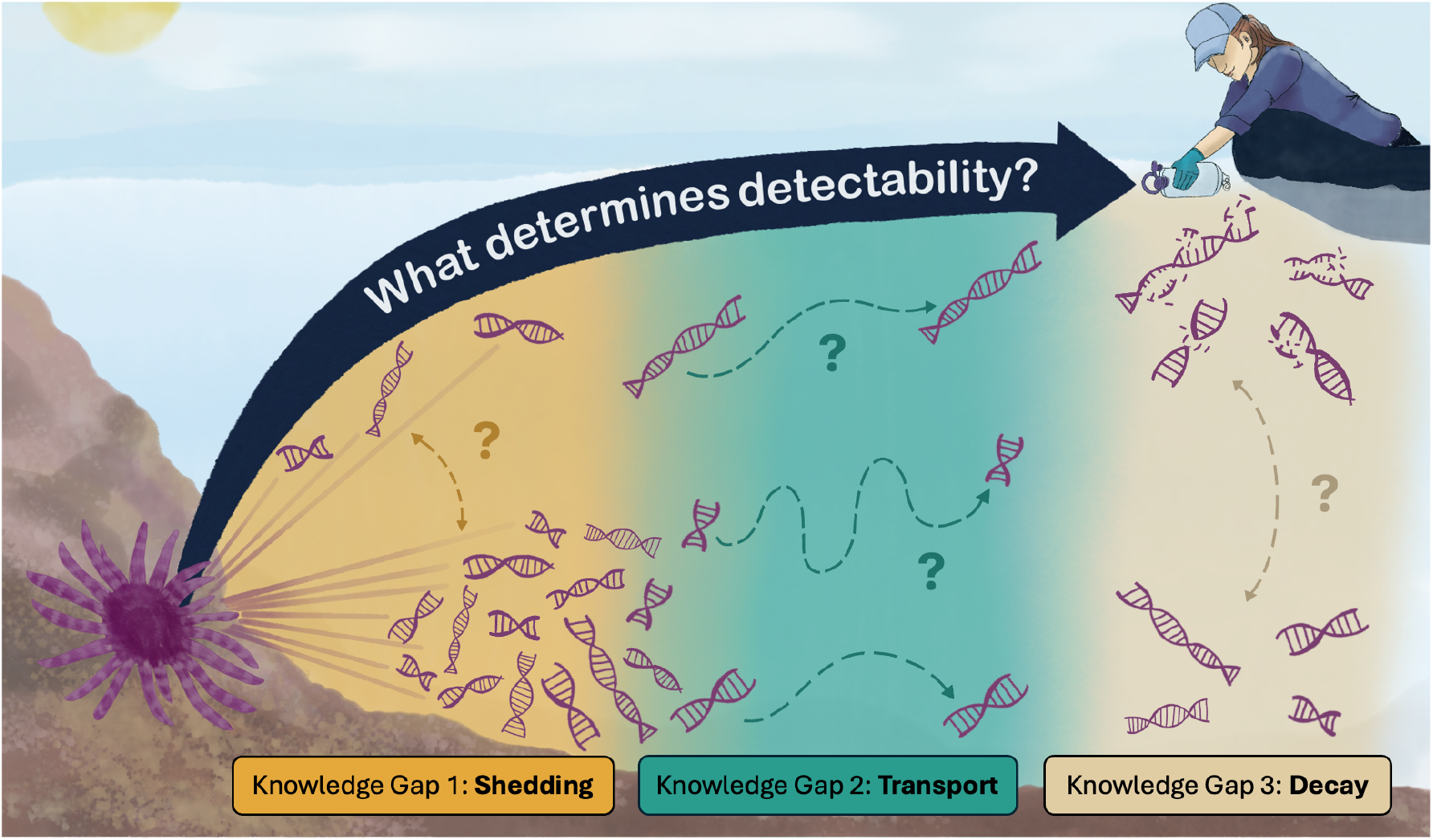
Conceptual model illustrating the processes that govern concentrations of eDNA from an organism, like *P. helianthoides*, in the environment

This study aims to (a) quantify eDNA shedding rates for *P. helianthoides* individuals across a range of sizes while monitoring organism stress levels, (b) quantify and compare decay rate constants for *P. helianthoides* eDNA under different sunlight conditions, and (c) contextualize the shedding and decay rates by conducting a systematic search and review of literature reporting marine eDNA shedding rates and decay rate constants. Thus, our work contributes to addressing shedding and decay knowledge gaps, as illustrated in Figure 1, and can underpin future work on eDNA fate in the environment. Specifically, quantitative measures of *P. helianthoides* shedding and decay can assist in the interpretation of measured eDNA concentrations in the coastal ocean when integrated into a process-based model of eDNA fate and transport. This work is especially important given that shedding and decay dynamics for the phylum *Echinodermata* are relatively understudied among marine organisms. More broadly, this study provides a vital foundation for the development of eDNA monitoring approaches for other organisms, environments, and contexts. Our shedding and decay experiments, with our detailed methodological reporting, could be replicated alongside the development of future eDNA assays, providing crucial information for both validating eDNA assays and contextualizing eDNA samples. Additionally, if context-specific shedding and decay experiments are not possible for a given monitoring program, our systematic search and review can help practitioners estimate the shedding rates and decay rate constant that might be applicable in their system.

## Materials and Methods

### Reproducibility%

To improve reproducibility, we have followed a variety of relevant reporting guidelines, including the Environmental Microbiology Minimum Information (EMMI) Guidelines^32^ and the Minimum Information for Publication of Digital PCR Experiments (dMIQE) Guidelines^33^. Completed EMMI and dMIQE checklists can be found in the Supporting Information (SI). We have also published detailed protocols for the methods described below including sample collection^34^, DNA extractions^35,36^ and ddPCR^37^, an RMarkdown file that fully reproduces the Results section of the paper (preliminary Zenodo archive available on request), and all raw data (preliminary Dryad archive available on request).

### Decay Experimental Design

The decay experiment was conducted outside of Sunflower Star Laboratory in Moss Landing, CA (36.80550° N, -121.78624° W). Two constant temperature water baths were set up approximately 1.5 meters from one another, one with exposure to incident sunlight (treatment) and one in an enclosed shed (dark control); to further shield the dark control from sunlight, a thick styrofoam lid was utilized. Water baths were circulated to keep water temperatures consistent throughout the experiment; temperatures were monitored using an infrared thermometer (Fisherbrand™, Thermo Fisher Scientific, Waltham, MA, USA) at least three times per day for the duration of the experiment.

At the start of the experiment, we filled 100 single-use 1000 mL enteral feeding pouches (Covidien, Dublin, Ireland) with seawater containing *P. helianthoides* eDNA. The eDNA-containing seawater was obtained from a recirculating aquaculture system housing 36 juvenile *P. helianthoides* individuals (ranging in size from 5-15 cm in diameter). The seawater used to fill the recirculating system was obtained from an on-site holding tank which was filled with seawater collected from an offshore intake in Monterey Bay, CA, 0.5 µm pore-size filtered, and UVC sterilized, limiting and changing background concentrations of eDNA and other constituents. The *P. helianthoides* eDNA in the seawater was shed naturally by the captive organisms and thus is a good representation of the eDNA that is likely to exist in the actual environment. Pouches were randomly assigned to each of the two experimental conditions and placed in the water baths so that they floated on the surface of the water (Figure 1B). We also filled 9 enteral feeding pouches with seawater from the on-site seawater holding tank for use as field blanks, which were distributed in randomized locations across the two water tables.

Upon adding the pouches to the system (t = 0), 5 pouches were destructively sampled from the treatment and control for immediate processing; we also removed one field blank pouch at this time. We then destructively sampled 5 pouches from each experimental condition at the following time points: 1 hour, 3 hours, 6 hours, 9 hours, 12 hours, 24 hours, 48 hours, 72 hours, and 96 hours. At one time point per day, we also sampled two field blank pouches, one from each experimental condition.

Before the experiment, we used a SpectriLight ILT 950 fiber optic radiometer and SpectriLight III processing software (International Light, Peabody, MA, USA) to measure the irradiance through the enteral feeding pouch material (Figure S3). On the first day of the experiment, we collected hourly irradiance measurements from directly above the pouches in each water table from before dawn to after dusk using the radiometer; on subsequent days, we collected irradiance measurements four times per day. Daily average UVA and UVB irradiance was calculated using the irradiance measurements (see SI).

### Shedding Experimental Design

The shedding experiment was conducted at Sunflower Star Laboratory in Moss Landing, CA. Twelve lidded aquaria were filled with a 10% bleach solution overnight and then rinsed twice with sterilized seawater. Sterilized seawater was generated from natural seawater obtained from an offshore intake in Monterey Bay, CA; the seawater was 0.5 µm pore-size filtered and UVC sterilized. After the aquaria were cleaned, they were filled with 60 L of sterilized seawater, which is the typical medium used for keeping sunflower stars at Sunflower Star Laboratory. We placed the aquaria in two circulating water baths, so that a consistent temperature could be maintained in the aquaria. One air stone was placed in each aquaria to facilitate aeration and mixing (Figure 1A). A thirteenth aquaria was prepared identically and set-up adjacent to the water baths with no added star, for use as a mesocosm control.

In advance, 530 mL polypropylene sampling bottles (PolyLab, Rai, Haryana, India) and PVC pipes cut to an appropriate length for fully mixing the aquaria were sterilized by soaking in a 10% bleach solution followed by 3 rinses in deionized water. Sampling material sterilization was conducted at a separate location from the Sunflower Star Laboratory and no sampling materials were re-used during the experiment, to minimize the potential for contamination with *P. helianthoides* DNA.

Samples were first collected from all thirteen aquaria before any *P. helianthoides* individuals had been added (t = 0). To sample, we stirred each aquaria in a circular motion three times using a sterilized single-use PVC stirring stick before using gloved hands to fill the sampling bottle from the surface of the aquaria. For every sample collected, we also measured the water surface temperature using a Mini IR Traceable™ Thermometer (Fisherbrand™, Thermo Fisher Scientific, Waltham, MA, USA). The water in a given sampling bottle (530 mL) was then transferred into a single-use enteral feeding pouch (Covidien, Dublin, Ireland). After water samples were collected, the volume removed from each aquarium was replaced with sterilized sea water to maintain a constant volume.

After the initial samples, a *P. helianthoides* individual was placed into each of the twelve aquaria. The individuals were all members of a cohort of stars raised at the Sunflower Star Laboratory from a spawning event on 14 February 2024, but were intentionally selected to encompass a range of star size characteristics. From measurements taken at the conclusion of the experiment, the stars ranged in size from 2.8 cm to 18.6 cm (arm tip-to-tip) and in weight from 1.17 g to 192.05 g (Table 3). We subsequently collected water samples from each aquaria at the following time points following the addition of the stars: 1 hour, 24 hours, 48 hours, 66 hours, 69 hours, and 72 hours. At one sampling point per day, we also collected a water sample from the field control aquaria.

**Table 3:** Summary of shedding rates per individual (copies/hr/individual) and shedding rate per biomass (copies/hr/g) with their propagated standard deviation as well as biomass (g) and tip-to-tip arm length (mm) for each of the twelve *P. helianthoides* individuals included in our shedding study.

|  | Per individual<br>(copies/hr/individual) |  | Per biomass<br>(copies/hr/g) |  | Organism characteristics |  |  |
| --- | --- | --- | --- | --- | --- | --- | --- |
| Star | Shedding rate | Propagated standard deviation | Shedding rate | Propagated standard deviation | Biomass (g) | Tip-to-tip arm length (mm) | Composite stress score |
| 1 | 22,200 | 6,500 | 19,000 | 5,600 | 1.17 | 28 | 1 |
| 2 | 24,000 | 6,000 | 19,800 | 5,000 | 1.21 | 31 | 1 |
| 3 | 42,300 | 15,000 | 8,190 | 2,800 | 5.17 | 55 | 0 |
| 4 | 22,900 | 9,000 | 1,390 | 550 | 16.48 | 86 | 0 |
| 5 | 89,600 | 10,000 | 3,790 | 420 | 23.66 | 112 | 0 |
| 6 | 548,000 | 96,000 | 15,300 | 2,700 | 35.85 | 113 | 7 |
| 7 | 134,000 | 52,000 | 2,780 | 1,100 | 48.13 | 121 | 0 |
| 8 | 55,800 | 12,000 | 896 | 200 | 62.28 | 127 | 0 |
| 9 | 133,000 | 6,800 | 2,700 | 140 | 49.22 | 142 | 2 |
| 10 | 91,300 | 17,000 | 1,080 | 200 | 84.37 | 143 | 1 |
| 11 | 64,900 | 11,000 | 4,230 | 700 | 15.33 | 144 | 0 |
| 12 | 450,000 | 87,000 | 2,350 | 460 | 192.05 | 186 | 5 |

The stars were not fed for the duration of the shedding experiment, and health checks were conducted three times per day primarily to count arm curls, which are known indicators of stress. Health checks were scored from 1 to 5, with 1 indicating no stress response detected; if a star received a health check score of 3 or higher during two successive health checks, they would have been removed from the experiment. A composite stress score was calculated for each star by summing all health check values (n = 8) and subtracting the baseline value for no detected stress response (8: a score of 1 across all 8 health checks); an additional point was added to each score for each time it was noted that the star had a stress response when added or removed from the experiment. Given the conditions for removing a star from the experiment, the range of possible composite stress scores would be 0 (no stress response detected) to 12.

### Water Filtration

All samples in single-use enteral feeding pouches were processed following Shea and Boehm^38^ to capture eDNA on filters. In short, enteral feeding pouches were hung and a 0.22 μm pore size PVDF Sterivex cartridge (MilliporeSigma, Burlington, MA, USA) was attached to the tubing on the pouch using a sterilized luer lock connector; water passed through the filter under the force of gravity. We conducted filtration in the shade to prevent any degradation by sunlight^30^.

After samples finished filtering (approximately 2-4 hours), the Sterivex cartridges were dried by passing air through them using a sterile syringe, secured with sterilized caps, placed in a small Whirl-Pak bag (Whirl-Pak, Madison, WI, USA), and transferred immediately to a -20°C freezer. Within 5 days, samples were transported on ice from the Sunflower Star Laboratory to Stanford University for DNA extractions where they were stored at -20°C.

### DNA Extraction and Quantification

We extracted DNA from the Sterivex cartridges within 4 months of receipt at the laboratory using the DNeasy Blood and Tissue Kit (Qiagen, Germantown, MD, USA) and modifications first detailed in Spens et al.^39^, as described by Shea and Boehm^35^. In short, we first added proteinase K and ATL buffer directly into the Sterivex cartridge and incubated for 24 hours at 56°C. After incubation, we removed and measured the liquid from each cartridge using a syringe and mixed it with equal volumes of AL buffer and 0°C molecular-grade ethanol. From there, we followed the manufacturer’s protocol for the remainder of the extraction; DNA was eluted in 100 µL of Buffer AE (Qiagen, Germantown, MD, USA). All DNA extractions were performed in a biosafety cabinet in a separate room from all post-extraction processing. In each batch of extractions, we included one negative extraction control where the initial sample volume was replaced with DNA-grade water (Fisher Scientific, Pittsburgh, PA, USA). All extracts were kept on ice until used as template in PCR as described below, typically within 2 hours (PCR was run directly after DNA extraction).

Droplet digital PCR (ddPCR) was used to quantify copies of *P. helianthoides* DNA in each sample. Undiluted DNA extracts were used as template. All ddPCR processing was performed using the AutoDG Automated Droplet Generator (Bio-Rad, Hercules, CA, USA), a C1000 Touch Thermal Cycler (Bio-Rad, Hercules, CA, USA), and a QX200 Droplet Digital PCR System (Bio-Rad, Hercules, CA, USA). Nucleic-acids were amplified using a species-specific hydrolysis probe assay for *P. helianthoides* targeting a 123 base pair fragment of the NADH dehydrogenase 5 (*nad5*) gene, located on the mitogenome (Table 1)^22^. The 20 µl reaction mixture consisted of 5 µL template DNA, 1 µL of a primer and probe pre-mix (900 nM forward and reverse primers and 250 nM probe; IDT, Coralville, IA, USA), 10 µL Bio-Rad ddPCR Supermix for probes, no dUTP (Bio-Rad, Hercules, CA, USA), and 4 µL nuclease-free water. Thermal conditions for the ddPCR amplifications were an initial denaturation for 10 minutes at 95°C followed by 45 cycles of 15 seconds at 95°C and 60 seconds at 60°C. We have published a full ddPCR protocol^37^ utilizing the Better Biomolecular Ocean Practices Template (BeBOP) template^40^.

**Table 1:**
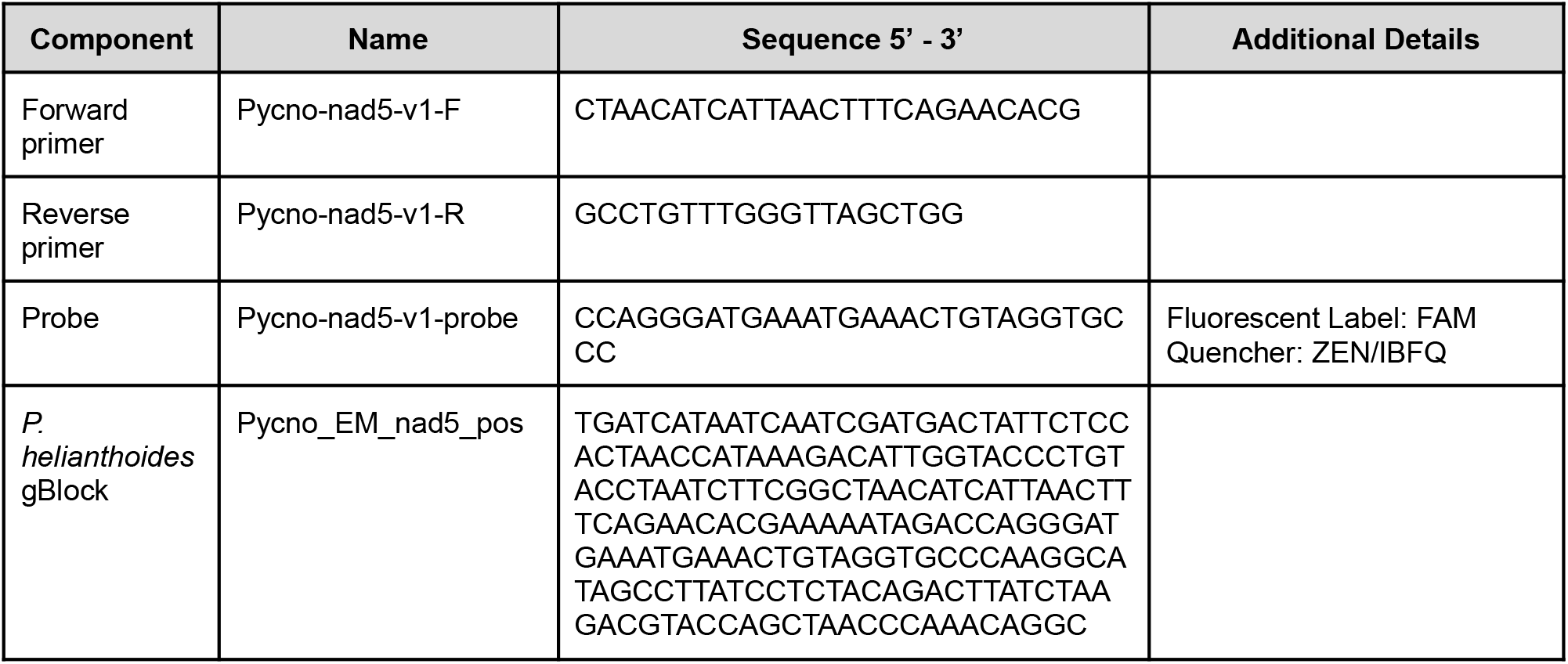
Sequence information for primers, probe, and positive control gBlock for *P. helianthoides* species-specific assay^22^.

Nucleic-acids were amplified in duplicate, and each 96 well plate processed contained one no-template control (NTC) consisting of nuclease-free water in place of template DNA and one positive control using a *P. helianthoides* gBlock (Table 1; IDT, Coralville, IA, USA) at a template concentration of ∼50 copies per µL template. Because we quantified samples immediately after extractions, we typically worked in batches of 48 samples (half plate). After quantification, technical duplicates were merged and a threshold was manually set above any variation in the cloud of negative droplets in the NTC for each plate using QX Manager (version 2.2.0; Bio-Rad, Hercules, CA, USA). If a replicate did not contain sufficient (>10,000) generated droplets, the value from the remaining replicate was used in place of a merged value. In order to be scored as positive, a sample had to have 3 positive droplets across the merged wells.

Any sample with 1 or 2 positive droplets was scored as negative. We further compared all samples to limits of detection and limits of quantification calculated in our laboratory during the development of the assay^22^. We conducted inhibition testing on a representative subset of five high concentration nucleic-acid extracts obtained from samples of the shedding and decay experiments. Nucleic-acids were used as template in ddPCR assays neat and diluted 1:5 and 1:10 in nuclease free water. The resultant concentrations were compared, after adjusting for dilution to determine if there was evidence of inhibition^41^.

### Data Analysis

All data analysis was conducted in R (version 4.4.2). An RMarkdown file that fully reproduces the results section of this paper is archived on Zenodo (Anonymized Reviewer Link: https://anonymous.4open.science/r/pycno-shedding-decay-peerreview-anonymized/).

#### ddPCR measurements

The ddPCR instrument provides the concentration of targets in units of copies per µL of the reaction mix along with 68% confidence intervals. We converted these values into values with units of copies per L of the original water sampled (C_x_) using the following equation (Equation 1):

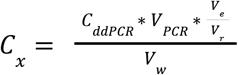

where C_ddPCR_ is the concentration generated by the ddPCR droplet reader (copies per µL in the reaction mix), V_PCR_ is the volume of the total PCR reaction (µL), V_e_ is the total volume post DNA extraction (µL), V_r_ is the volume of template added to the PCR reaction (µL), and V_w_ is the volume of sample water filtered onto the Sterivex for extraction (L).

#### Decay rate constant calculations

First order decay rate constants (k) with units per hour were calculated assuming decay of the target is first order. They were derived by plotting ln(C/C_0_) versus time and applying a linear model to obtain k and standard error (SE) using the *lm* function in R, where C is the eDNA concentration (copies/L) and C_0_ is the average eDNA concentration at the start of the experiment in each condition (copies/L). The y-intercept was set to 0.

To test the null hypothesis that the decay rate constant for the samples in the treatment (light) is the same as the decay rate constant for the samples in the control (dark), we performed a z-test using the following equation (Equation 2), following Andruszkiewicz et al.^30^:

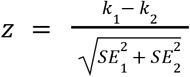

To reject the null hypothesis at α = 0.05, |z| must be greater than 1.96.

#### Shedding rate calculations

To calculate eDNA shedding rates, we modeled the concentration of sunflower star eDNA in the shedding experiment aquaria assuming the aquaria were completely mixed batch reactors^25,42^ using the following equation (Equation 3):

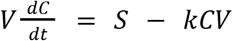

where V is the volume of the aquaria (L), C is the concentration of eDNA (copies/L), t is the time elapsed (hr), S is the eDNA shedding rate (copies/hr), and k is the decay rate constant (1/hr). At steady state (Equation 4):

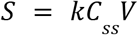

where C_ss_ is the steady-state eDNA concentration (copies/L), which for this experiment was defined as the average eDNA concentration across the final three time points (t = 66, 69, 72 hours). V was considered a constant, as we determined that the sampling with replacement that we conducted did not impact modeling results (see SI).The time to steady state in this system assuming a k value on the order of those observed in other studies (∼0.1 hour^-1^) would be approximately 50 hours (see SI). Following Sansom & Sassoubre^43^, to determine the error associated with the calculated shedding rate, we propagated the errors associated with k and C_ss_ using the following equation (Equation 5):

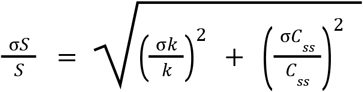

To facilitate comparisons across aquaria and with other studies, we also calculated biomass-normalized shedding rates.

To determine whether shedding rates varied significantly among star individuals, we used the *lme4* package in R (version 2.0.1)^44^ to fit random-intercept linear mixed-effects models with individual star as a random effect for both shedding rate per individual and per biomass (shedding rate ∼ 1 + (1 | star identity)). The models were fit to all steady-state time points (t = 66, 69, 72 hours) so that we could assess both between-individual and within-individual variance. We determined whether the random intercept was statistically significant using a likelihood ratio test which compared the full model to a model version without the random effector, using the *lmerTest* package (version 3.2.1)^45^.

To determine whether organisms’ characteristics predicted variation in shedding rates, we fit multiple linear regression models with log-transformed shedding rates (per individual and per biomass) as the response variable and biomass, tip-to-tip arm length, and stress score as predictors (log(shedding rate) ∼ biomass + tip-to-tip arm length + composite stress score).

These models were fit to average shedding rate values, rather than all steady-state time points, to avoid pseudo-replication. We used variance inflation factors and pairwise Pearson correlations to assess predictor collinearity.

### Literature Review

To compare our calculated shedding and decay rates with other marine organisms, we conducted a systematic search and review^46^ of existing peer-reviewed literature. We searched the title and abstract of all papers indexed by Web of Science, Scopus, and PubMed for the following search terms: *[eDNA OR “environmental DNA”]*, *[rate*]*, and *[shed* OR decay* OR degrad* OR releas*]*. We supplemented these identified papers with all literature included in several previous reviews of eDNA shedding and decay rates^24–26,47–49^. From these papers, we included in our review any study that collected eDNA samples from marine waters (natural or artificial) and reported either monophasic decay rate constants (in units of hr^-1^ or day^-1^) or shedding rates (in units of pg/hr/individual, pg/hr/g, copies/hr/individual, or copies/hr/g).

## Results

### ddPCR Assay Quality Control

Procedurally, we followed a limit of detection of 3 positive droplets per merged replicates, which corresponds to approximately 0.04 copies/L for the decay experiment and 0.08 copies/L for the shedding experiment. From previous assay validation work, we also calculated limits of detection and limits of quantification using multiple methods and scenarios^22^. Applied to this study, the most conservative limit of detection and limit of quantification, derived using the Klymus et al.^50^ method and also requiring 3 positive droplets to constitute a detection, would be 194 copies/L (LOD) and 480 copies/L (LOQ) for the decay experiment and 366 copies/L (LOD) and 906 copies/L (LOQ) for the shedding experiment. All positive controls and experimental samples (excluding t=0 shedding samples), were above even this conservative limit of quantification. Concentrations obtained from the 5 samples tested for inhibition indicated no inhibition (Figure S6).

Across both the shedding and decay studies, we processed 9 no-template controls (NTCs, no amplification), 9 positive controls (all amplified as expected), 9 extraction blanks (all below the limit of detection), 13 field blanks (all below the limit of detection), 84 experimental shedding samples, and 100 experimental decay samples. 7 of 224 (3%) samples had one ddPCR technical replicate without sufficient droplets generated; the remaining ddPCR replicate was used in lieu of merging the technical replicates. 31 samples (15%) were re-run after one freeze-thaw due to errors with the ddPCR instrument.

### eDNA Decay

Across the experiment, average temperature in the treatment (light) was 15.6°C ± 1.9°C and in the control (dark) was 17.6°C ± 1.7°C. Temperatures were elevated in the dark treatment due to water pumping and cooling machinery housed in the same shed, which produced waste heat. After correcting for losses from the pouch material and screening by the water, the experiments were exposed to daily average UVA irradiance of 3.76 W/m^2^ in the treatment (light) and 0.03 W/m^2^ in the control (dark); and daily average UVB irradiance of 0.09 W/m^2^ in the treatment (light) and 0.01 W/m^2^ in the control (dark).

Both the treatment (light) and control (dark) exhibited patterns of first order decay (Table 2; Figure 3A); all adjusted R^2^ values were above 0.85. In the dark control, *P. helianthoides* eDNA had a decay rate constant of 0.0270 ± 0.0016 hr^-1^; in the light treatment, *P. helianthoides* eDNA had a decay rate constant of 0.0244 ± 0.0011 hr^-1^ (Table 2; Figure 3B). Using a z-test with α = 0.05 (|z| = 1.39), we did not reject the null hypothesis that the treatment and control have the same decay rate constant. Therefore, we calculated a combined decay rate constant across all samples (0.0257 ± 0.0010 hr^-1^), which we use as the k value for subsequent analyses.

**Figure 2:**
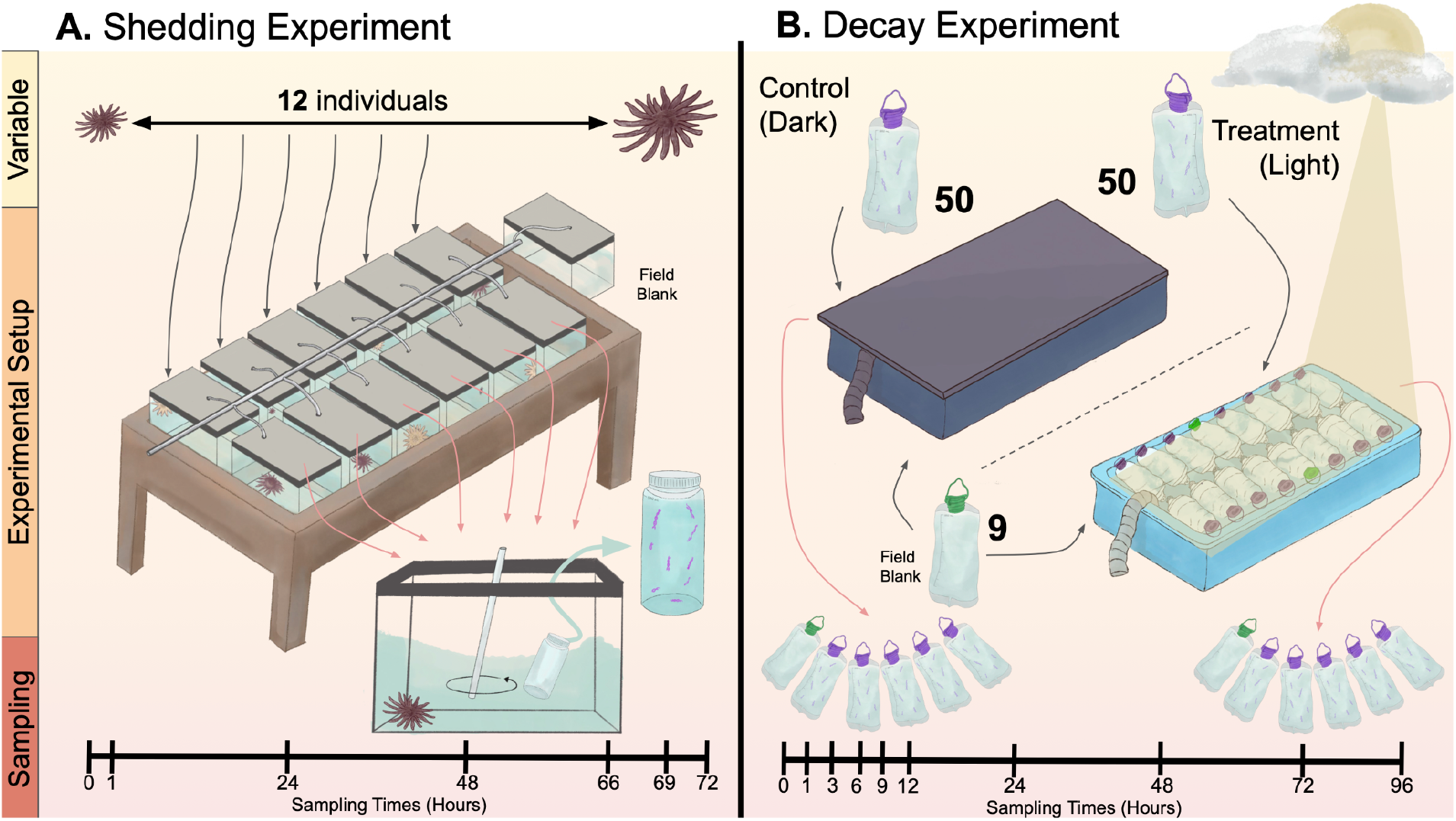
Experimental set-up for shedding study (A) and decay study (B)

**Figure 3:**
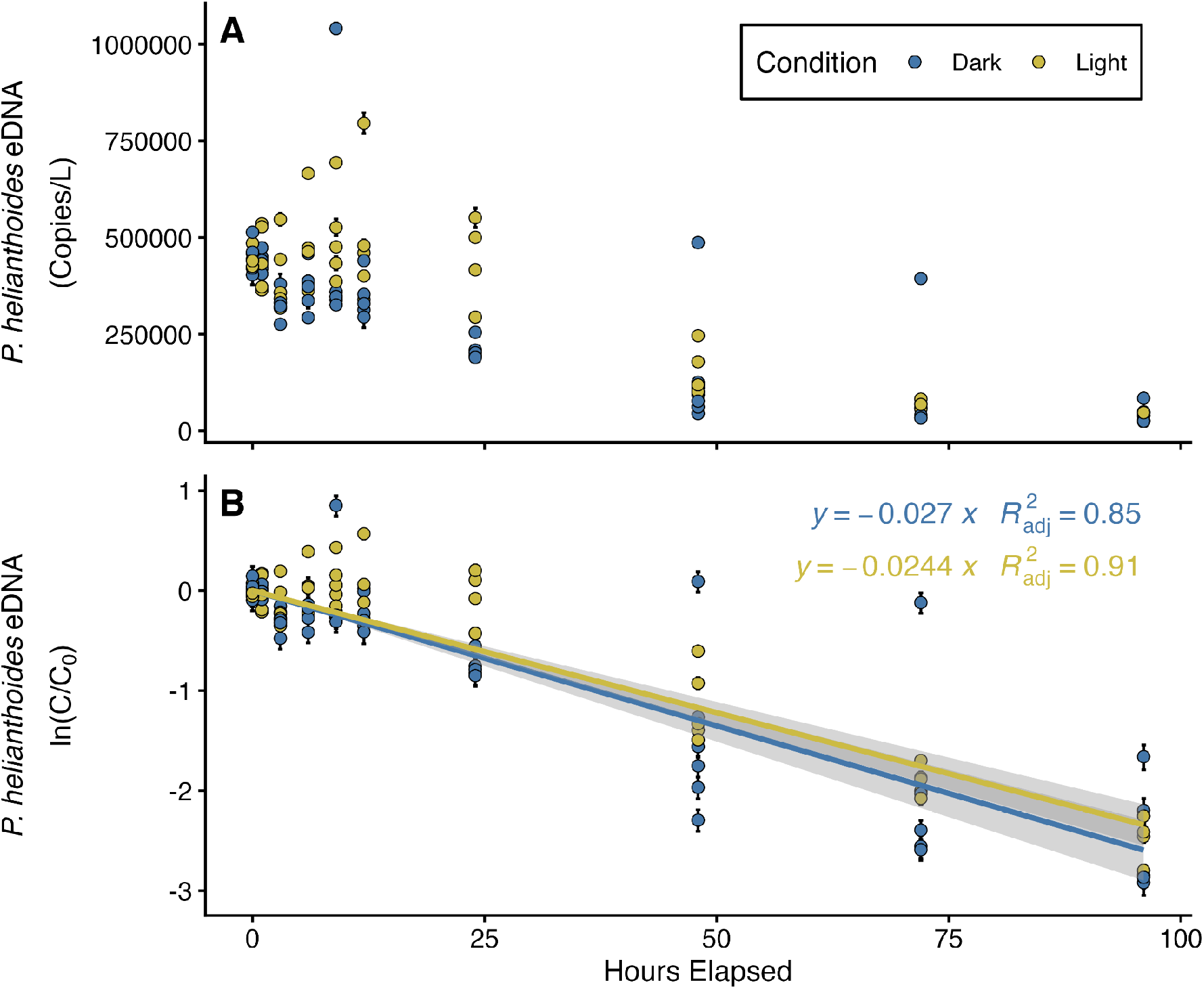
eDNA concentrations (A) and the natural log of normalized eDNA concentrations (B) over time for treatment (light) and control (dark). Each point represents merged technical replicates. In Panel A, error bars represent 95% total error (incorporating Poisson error and variance among replicates). In Panel B, error bars represent the propagated standard deviation of error from C and C_0_ values.

**Table 2:** Summary of decay rate constants (hr^-1^), their standard error (SE), the adjusted R^2^ of the first order decay model, and the calculated half-life of *P. helianthoides* eDNA in each of our experimental conditions, as well as combined across all samples.

| Condition | First order decay model $k$ ($\text{hr}^{-1}$) | SE | Adjusted $R^2$ | Half-life $[\ln(2)/k]$ (hr) |
| --- | --- | --- | --- | --- |
| Control (Dark) | 0.0270 | 0.0016 | 0.852 | 25.6 |
| Treatment (Light) | 0.0244 | 0.0011 | 0.914 | 28.4 |
| All Combined | 0.0257 | 0.0010 | 0.878 | 26.9 |

### eDNA Shedding Rates

Across the shedding experiment, all aquaria with stars were maintained at an average temperature of 12.9°C ± 0.5°C. The field control aquaria, which was not in a water bath to maximize space for experimental aquaria, was maintained at an average temperature of 18.1°C ± 1.2°C. At t = 0, no water samples from any aquaria were above the limit of detection, confirming that there was no background *P. helianthoides* eDNA signal in the aquaria before organisms were added. After organisms were added to aquaria, eDNA concentrations and assessed stress responses varied across the experiment (Figure 4). Composite stress scores for each organism, which were calculated by summing scores from health checks and stress responses on addition or removal from the aquaria, ranged from 0 to 7.

**Figure 4:**
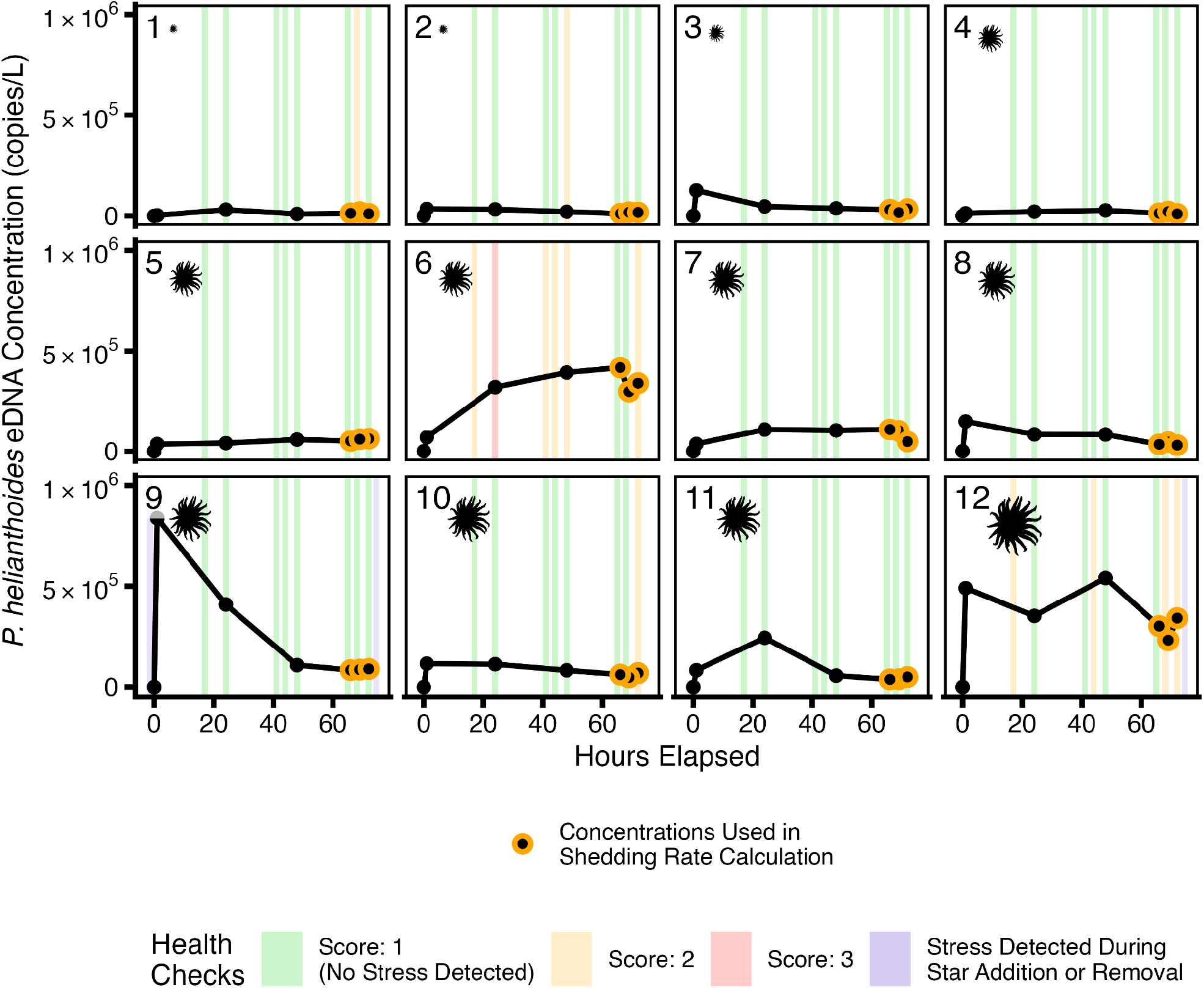
*P. helianthoides* eDNA concentrations (copies/L) measured for each organism over the duration of the experiment. Highlights show the results from health checks conducted throughout the experiment to assess star stress. The final three time points, later used in shedding rate calculations, are outlined in yellow.

Calculated shedding rates were variable across our 12 study organisms (Table 3; Figure 5), as assessed with a random-intercept fixed-effects model with individual organisms as a random effect. Through a likelihood ratio test, we confirmed that between-individual variation was significant (◻^2^ = 56.93, df = 1, p < 0.001). Shedding rates per individual ranged from 22,200 to 548,000 copies/hr/individual (average: 140,000 ± 170,000). Shedding rates normalized by biomass ranged from 896 to 19,800 copies/hr/g (average: 6,790 ± 7,100).

**Figure 5:**
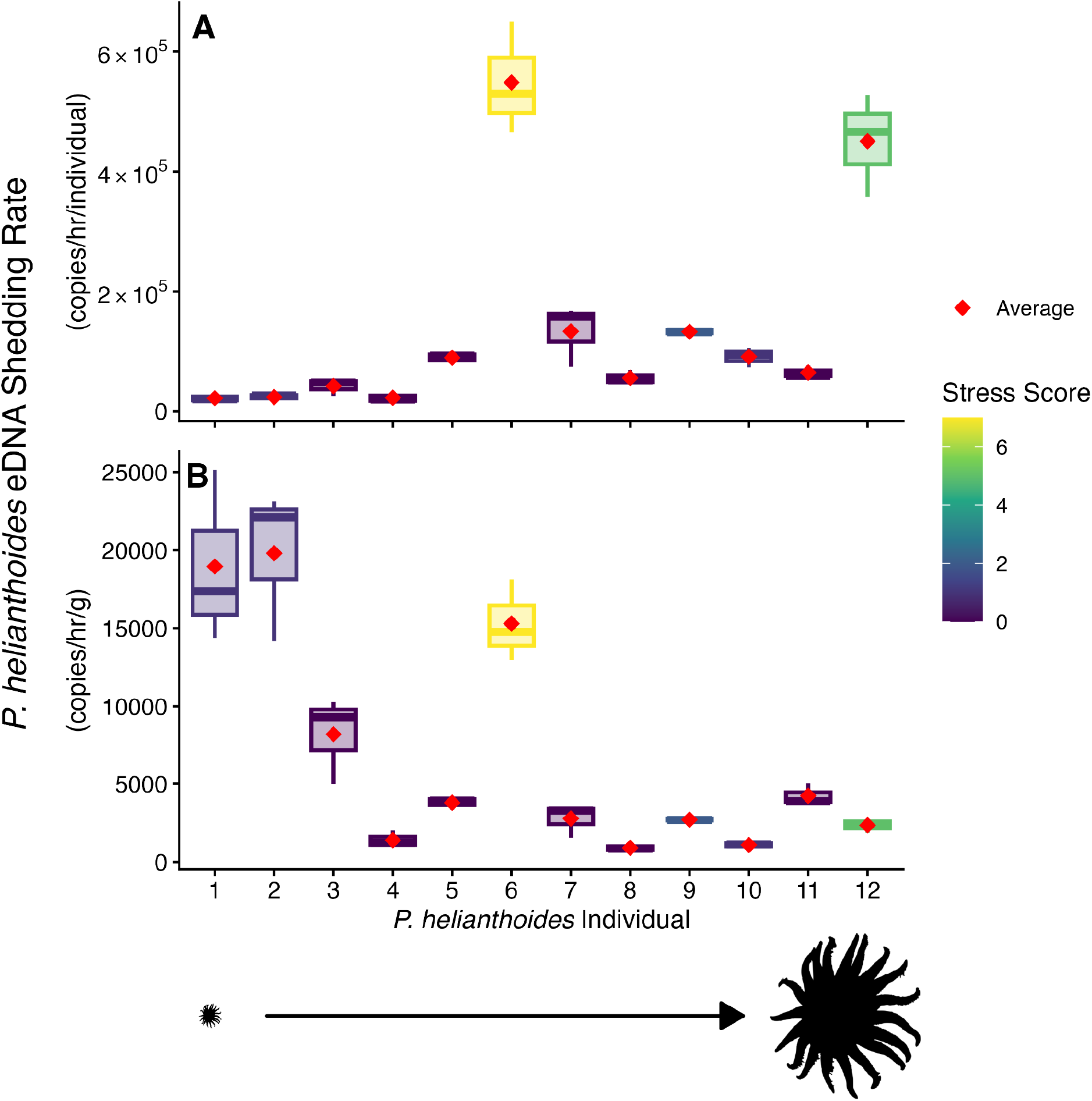
*P. helianthoides* eDNA shedding rates normalized by individual (A) and normalized by biomass (B) for each of twelve individual organisms in our shedding study. Box plots depict the range of shedding rates calculated from individual concentration values across our final three time points (66, 69, and 72 hours); red diamonds represent the shedding rates calculated from the average concentration values across our final three time points. Individual stars are organized from smallest tip-to-tip arm length (1) to largest tip-to-tip arm length (12). The color of the box plots corresponds to a composite stress score, which combines health check scores and any stress responses noted during star addition or removal.

To better understand differences across organisms, we used multiple linear regression models to determine if organism characteristics explained the variance in both log-transformed shedding rates per individual and per biomass. We found that models explained a substantial proportion of the variance for both measures, suggesting that biomass, tip-to-tip arm length, and composite stress score together accounted for much of the observed individual variation in shedding (shedding rate per individual: adjusted R^2^ = 0.8291; F(3,8) = 18.79, p = 0.001; shedding rate per biomass: adjusted R^2^ = 0.65; F(3,8) = 7.83, p = 0.009).

Across both models, the strongest organism characteristic was composite stress score, which was a significant predictor even when accounting for other characteristics (shedding rate per individual: β = 0.303, p = 0.002; shedding rate per biomass: β = 0.299, p = 0.016). Tip-to-tip arm length was also a significant predictor of shedding rate per individual (β = 0.014, p = 0.010) but not a significant predictor of shedding rate per biomass (β = -0.013, p = 0.064). Biomass was not a significant predictor in either model (shedding rate per individual: p = 0.492; shedding rate per biomass: p = 0.320). While tip-to-tip arm length and biomass were moderately correlated (r = 0.77), variance inflation factors were low for all predictors (biomass: 2.93; tip-to-tip arm length: 2.47; composite stress score: 1.32), which suggests that multicollinearity did not substantially influence the models.

### Systematic Search and Review

The systematic search and review identified 29 total papers: all papers reported marine eDNA decay rate constants (in the units hr^-1^ or day^-1^) and 9 papers (31%; n = 29) also included shedding rates (in the units pg/hr/individual, pg/hr/g, copies/hr/individual, or copies/hr/g) (Table 4). Across all papers, we extracted 134 individual decay rate constants and 73 individual shedding rates, across a range of organism types, water temperatures, and experimental conditions (Figure 6). The full summary of decay rates constants and shedding rates can be found in the SI.

**Figure 6:**
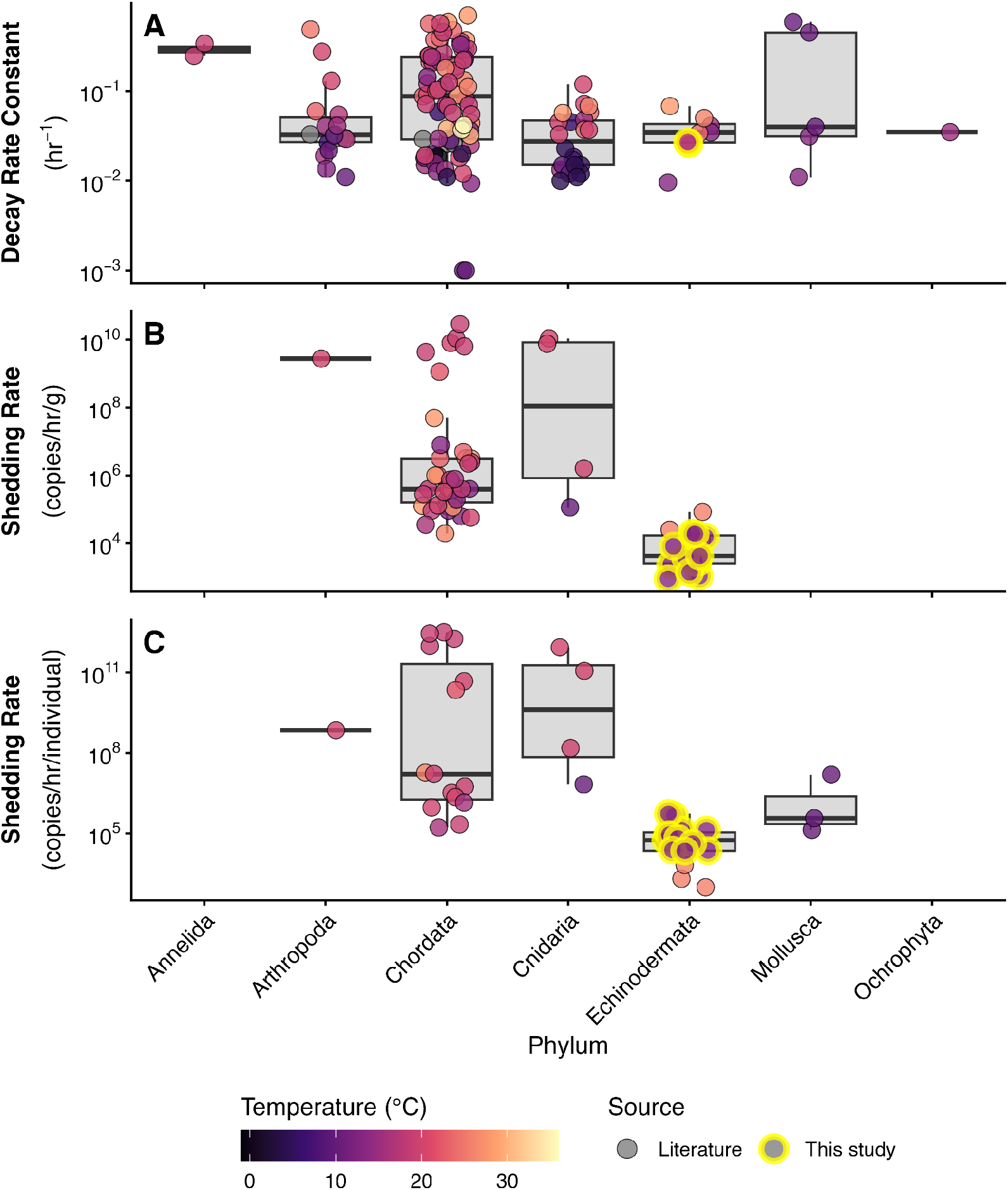
Summary of eDNA decay rate constants (A), shedding rates in copies/hr/g (B), and shedding rates in copies/hr/individual (C) found in our systematic review as a function of phylum. The shading of the points depicts the temperature of the shedding or decay experiment (when specified); points outlined in yellow were calculated in this study. Three Chordata decay rate constants, derived from experiments at pH = 4, were reported as negative values; these are not depicted in panel A due to their incompatibility with the log scale.

**Table 4:** Summary of papers identified in our systematic review that report marine eDNA decay rate constants or shedding rates.

| Study | Number of Values Reported |  | Organisms | Variables Tested |
| --- | --- | --- | --- | --- |
|  | Decay Rate Constants | Shedding Rates |  |  |
| Andruszkiewicz Allan et al., 2021<sup>25</sup> | 15 | 8 | *Fundulus heteroclitus*, *Palaeomenes* spp., *Aurelia aurita*, *Chrysaora* spp. | Organism, Temperature, Sunlight |
| Andruszkiewicz et al., 2017<sup>30</sup> | 2 | None | *Scomber japonicus* | Sunlight |
| Brasseale et al., 2025<sup>51</sup> | 1 | None | *Tursiops truncatus* | NA |
| Chung et al., 2026<sup>52</sup> | 1 | 2 | *Epinephelus akaara* | NA |
| Collins et al., 2018<sup>26</sup> | 16 | None | *Carcinus maenas*, *Lipophrys pholis* | Organism, Season, Inshore/Offshore |
| Cowart et al., 2018<sup>53</sup> | 1 | None | *Chionodraco rastrospinosus* | NA |
| Dass et al., 2025<sup>54</sup> | 1 | None | *Homo sapiens* | NA |
| Ellis et al., 2022<sup>55</sup> | 2 | None | *Undaria pinnatifida*, *Asterias amurensis* | Organism |
| Forsstrom & Vasemagi, | 1 | None | *Rhithropanopeus harrisii* | NA |
| 2016<sup>56</sup> |  |  |  |  |
| Holman et al., 2022<sup>57</sup> | 8 | None | *Asterias rubens*, *Ostrea edulis*, *Ciona intestinalis*, *Monocorophium insidiosum*, *Diplosoma spp.*, *Crepidula fornicata*, *Botrylloides violaceus*, *Amphipholis squamata* | Organism |
| Jo et al., 2017<sup>28</sup> | 2 | None | *Trachurus japonicus* | DNA Fragment Length |
| Jo et al., 2019<sup>58</sup> | 12 | 12 | *Trachurus japonicus* | Temperature, Biomass |
| Jo et al., 2020<sup>59</sup> | 12 | 12 | *Trachurus japonicus* | Temperature, Biomass, DNA Fragment Length |
| Jo et al., 2023<sup>60</sup> | 9 | None | *Danio rerio* | Temperature, pH |
| Kirtane et al., 2021<sup>61</sup> | 8 | 16 | *Centropristis striata*, *Pseudopleuronectes americanus*, *Paralichthys dentatus* | Organism, DNA Fragment Length |
| Kutti et al., 2020<sup>62</sup> | 1 | 2 | *Lophelia pertusa* | NA |
| Kwong et al., 2021<sup>63</sup> | 3 | 6 | *Acanthaster cf. solaris* | Temperature, Feeding |
| Lopez et al., 2025<sup>64</sup> | 3 | 3 | *Magallana [Crassostrea] gigas* | pCO<sub>2</sub> |
| McCartin et al., 2022<sup>65</sup> | 11 | None | *Desmophyllum pertusum* | Temperature, Dissolved Oxygen, pH |
| Minamoto et al., 2017<sup>66</sup> | 1 | 2 | *Chrysaora pacifica* | NA |
| Morrissey et al., 2022<sup>67</sup> | 2 | None | *Chironex fleckeri* | Temperature |
| Qian et al., 2022<sup>68</sup> | 4 | None | *Fenneropenaeus chinensis* | Temperature |
| Saito & Doi, 2021<sup>69</sup> | 2 | None | *Trachurus japonicus*, *Oncorhynchus kisutch* | Organism |
| Sassoubre et al., 2016<sup>42</sup> | 5 | 10 | *Engraulis mordax*, *Sardinops sagax*, *Scomber japonicus*, | Organism, Organism Combination |
| Sigsgaard et al., 2017<sup>31</sup> | 2 | None | *Rhincodon typus* | Sunlight |
| Skinner et al., 2025<sup>70</sup> | 1 | None | *Caretta caretta* | NA |
| Thomsen et al., 2012<sup>71</sup> | 2 | None | *Platichthys flesus*, *Gasterosteus aculeatus* | Organism |
| Weltz et al., 2017<sup>72</sup> | 2 | None | *Zearaja maugeana* | Dissolved Oxygen, Biomass |
| Wood et al., 2020<sup>73</sup> | 4 | None | *Sabella spallanzanii*, *Styela clava* | Organism, Organism Combination |
| This Study | 2 | 24 | *Pycnopodia helianthoides* | Sunlight |

In the literature, decay rate constants ranged from -0.007 to 0.696 hr^-1^ (n = 134; average = 0.121 hr^-1^). Shedding rates ranged from 1.28 x 10^4^ to 2.92 x 10^10^ copies/hr/g (n = 47; average = 1.73 x 10^9^ copies/hr/g) and from 988 to 3.19 x 10^12^ copies/hr/individual (n = 26; average = 3.82 x 10^11^ copies/hr/individual). In the phylum *Echinodermata,* decay rate constants ranged from 0.010 to 0.068 hr^-1^ (n = 6; average = 0.040 hr^-1^) and shedding rates ranged from 1.28 x 10^4^ to 8.40 x 10^4^ copies/hr/g (n = 3; average = 4.06 x 10^4^ copies/hr/g) and from 988 to 6,642 copies/hr/individual (n = 3; average = 3226 copies/hr/individual). As shown in Figure 4, our reported decay rate constants and shedding rates were among the lowest reported in studies of marine species, and on par with other studied echinoderms: *Acanthaster cf. solaris* (shedding and decay)^63^, *Asterias rubens* (decay only)^57^, *Asterias amurensis* (decay only)^55^, and *Amphipolis squamata* (decay only)^57^.

## Discussion

With the development of a *P. helianthoides*-specific eDNA assay^22^, there has been growing interest in monitoring *P. helianthoides* populations using coastal water samples. However, properly interpreting what eDNA concentrations measured in these samples tell us about the presence and abundance of *P. helianthoides* organisms requires quantitative information on shedding and fate of *P. helianthoides* eDNA. In this study, we provide first estimates of *P. helianthoides* eDNA shedding rates and decay rate constants, and among the first measurements for *Echinodermata* more broadly.

### eDNA Decay

*P. helianthoides* eDNA decay rate constant is 0.0257 hr^-1^ in sunlit and dark conditions. Through our systematic search and review, we found that our decay rate constant is within the lowest third of those reported in the literature across other studies of marine species, and similar to that of other echinoderms, including *Acanthaster cf. solaris* (k = 0.034 - 0.068 hr^-1^)^63^, *Asterias rubens* (k = 0.035 hr^-1^)^57^, *Asterias amurensis* (k = 0.041 hr^-1^)^55^, and *Amphipolis squamata* (k = 0.010 hr^-1^)^57^. Decay rate constants may be affected by temperature^25^, other physiochemical conditions^65^, microbial processes^74,75^, DNA fragment length^28^, and its form in the environment (mitochondrial, nuclear, dissolved, adsorbed to particles)^76^.

We did not find a difference between decay rate constants for our treatment (light) and control (dark) conditions, which is consistent with two other studies that have investigated the influence of sunlight on eDNA degradation in marine waters^30,31^. eDNA can be both broken down and altered by direct and indirect photochemical reactions, the influence of which can be modulated by the sequence of the DNA, the prevalence of light-absorbing dissolved organic matter, environmental conditions, and microbial communities. While research has confirmed many organic biomolecules, like DNA, are subject to photolysis by sunlight, continued research is necessary to disentangle the role of sunlight in eDNA degradation, specifically^29^. We note several experimental factors that likely affect the finding that sunlight does not affect *k*. For one, cloud cover reduced the measured UVA and UVB irradiance on three of the four days of our decay experiment (Figure S2). While the experimental irradiance was similar to another study conducted at a nearby location^30^, we may have observed a difference between our treatment and control conditions had there been less cloudy conditions during the experiment.

Additionally, the average temperature in the control water bath was elevated about 2°C from the treatment water bath, due to heat given off by water pumping and cooling machinery housed in the same shed as the control water bath. This means that the influence of sunlight could have been masked by the influence of temperature; that is, the potential increase in decay rate constant in the control samples due to higher temperatures may have been larger than the hypothesized increase in decay rate constant in the treatment samples due to sunlight exposure. However, another study of eDNA decay in marine waters across comparable small temperature differences did not find a significant effect of temperature on decay rate constants^63^. We also note that the ocean water used in the experiments was clear with very limited absorbance (Figure S4) suggesting limited presence of light absorbing dissolved organic matter. In the presence of sunlight, organic matter can sensitize production of reactive species such as singlet oxygen which may react and cleave DNA molecules^29^. Future work under more controlled conditions with a variety of waters will further elucidate the potential for sunlight to destroy eDNA biomonitoring targets.

Additionally, future work on the influence of temperature on *P. helianthoides* eDNA decay would be especially valuable given the reported influence of temperature on marine eDNA decay rates across our systematic search and review^24^. The large historic range of *P. helianthoides*–from Baja Sur California, Mexico (14 - 23°C) to the Aleutians, Alaska, USA (3 - 10°C)–means eDNA biomonitoring might occur over a wide range of oceanic conditions^77^.

Controlled laboratory experiments across a temperature range that corresponds with coastal conditions across the Northeast Pacific Ocean would complement the findings from our decay experiment.

### eDNA Shedding

Our calculated shedding rates ranged on the order of 10^4^ - 10^5^ copies/hr/individual and 10^3^ - 10^4^ copies/hr/g. The rates per individual were among the lowest reported in the literature for marine organisms, higher only than the rates reported for another juvenile echinoderm, *Acanthaster cf. solaris*^63^; rates normalized by biomass were the lowest reported in any study. This suggests that echinoderms may have different shedding mechanisms than other marine organisms, such as fish.

We note that our calculated biomass-normalized rates were also consistent with sampling in a flow-through system housing 39 adult *P. helianthoides* individuals. That study reports an average concentration of 1.78 x 10^6^ copies/L in samples from the mesocosm^22^. Using the methods described in Kwong et al.^63^, the flow rate of the system (3066 L/hr), and an estimated individual weight of 3 kg, shedding rate estimates from that study are approximately 10^8^ copies/hr/individual and 10^4^ copies/hr/g. These results align well with those reported in this study.

The low shedding rates may have behavioral causes. For one, *P. helianthoides* are relatively sedentary compared to many other marine organisms, especially when not feeding. This may comparatively limit the opportunities for cells to be physically sloughed into the water column. Additionally, our shedding experiment provides evidence that *P. helianthoides* individuals experiencing stress may shed more; composite stress score was the strongest statistically significant predictor of shedding rates, and specific noted stress behaviors often coincided with elevated eDNA concentrations throughout the experiment. It has been documented that mucus^78^ and saponin^79^ production can accompany stress responses in echinoderms; increased water pumping through respiration or metabolic processes could also contribute to elevated shedding. These secretions or metabolic processes may be important mechanisms for *P. helianthoides* eDNA release into the water. Continued research into *P. helianthoides* behavior, stress responses, and activity levels could help contextualize expectations for shedding rates in field conditions.

One unique feature of this experiment was the shared age, but varying size, of the individuals studied. Other studies of shedding rates across individuals necessarily have co-varying age and size^80^, making it challenging to disentangle the influence of each characteristic. In this study, all individuals studied were reared from the same spawning event, despite reaching quite different sizes by the start of our experiment; this enables us to isolate the influence of body size on shedding rates for the first time. Generally, we found that larger stars shed more per individual, but shed less per gram. This finding is consistent with several other studies, including with marine organisms. Maruyama et al.^80^ compared eDNA shedding of adult (large size) and juvenile (small size) bluegill sunfish and found that the eDNA release rate was higher for adult individuals, but the eDNA release rate per fish body weight was higher for juvenile individuals. This relationship is consistent with allometric scaling. However, the organisms used in our shedding experiment were still quite small compared to adult *P. helianthoides* in the wild. The largest organism in our study was 192 grams with a tip-to-tip arm length of 186 mm; in contrast, adult *P. helianthoides* are typically over 40 cm^81^. Given the variation in shedding rates we found across organisms of different sizes, it would be important to conduct future experiments with individuals more representative of the full size range of *P. helianthoides*, particularly larger individuals. The variation in shedding among individuals means that in exercises that link measured concentrations to organism abundance^82^ will need to consider a distribution of shedding rates in the model, representing the process as stochastic.

### Roadmap for Future Species-Specific Monitoring

For *P. helianthoides* this work advances the development of a large-scale eDNA monitoring program to map existing and recovering populations across the Northeast Pacific Ocean. While *P. helianthoides* have comparatively low eDNA shedding rates, significant research on the crown of thorns seastar (*Acanthaster cf. solaris*), the organism with the most similar shedding rates to *P. helianthoides* among those reported in the literature, has shown that they are detectable via a small number of eDNA samples at relatively low densities (3 individuals ha^-1^)^83^. This finding suggests that it will be possible to detect rare *P. helianthoides* individuals as well, given what we now know about their general shedding and decay dynamics as well as recent successful detections of *P. helianthoides* at low densities (< 1 individual per HA)^22^. Future modelling and field experiments could improve our understanding of *P. helianthoides* eDNA detection probabilities under realistic hydrodynamic conditions.

The high variability in shedding rates demonstrated in this study may complicate efforts to quantitatively interpret eDNA signals from coastal samples. We found that organism stress can influence eDNA shedding rates. A previous study also found elevated DNA concentrations in samples collected above a *P. helianthoides* individual showing symptoms of sea star wasting disease and from mesocosms containing P. helianthoides larvae^22^. Since the previous study did not enable shedding rate calculations, future work focused on characterizing shedding rates for *P. helianthoides* individuals across life history stages and health statuses could help constrain our understanding of what eDNA concentrations in environmental samples can tell us about organisms in the field. At the same time, the variability in shedding rates we found also reifies that eDNA monitoring should be one of many tools in a conservation toolkit, and that when possible, repeated sampling at the same locations is valuable.

As conservation practitioners and resource managers increasingly seek to monitor marine organisms using eDNA, the shedding and decay experiments conducted here can serve as a guide for others seeking to validate and contextualize new species-specific eDNA assays. Our experiments utilized existing aquaculture infrastructure, were able to be run simultaneously, and have been rigorously documented through published protocols. These features enable the experiments to be reproduced and adapted in other contexts. In scenarios where conducting new shedding and decay experiments is not feasible, the systematic search and review enables practitioners to identify a reasonable range of shedding rates and decay rate constants for their organism of interest. While our results demonstrate some of the many challenges in linking eDNA concentrations to abundance of organisms, they also showcase the promise of future conservation efforts using eDNA. Our experiments and systematic search and review have confirmed that *P. helianthoides* may be a particularly difficult organism to monitor via eDNA, given its comparatively low shedding rate, yet *P. helianthoides* have been successfully detected in field samples^22^. Further research into shedding and decay dynamics will continue to support the interpretation of eDNA biomonitoring data, continuing a growing and important tradition of applied conservation research using eDNA tools.

## Supporting Information

File containing additional information for dMIQE reporting guidelines (Figure S1), time to steady state derivation, and calculation of daily average UVA and UVB irradiance (Figures S2-S5; Table S1 (PDF); completed EMMI checklist (PDF); completed dMIQE checklist (PDF); outline of model to confirm the limited impact of sampling loss (Excel); systematic search and review shedding data (CSV); systematic search and review decay data (CSV)

## Supporting information

Supporting Information 1 - Additional Methodological Details

Supporting Information 2 - EMMI Checklist

Supporting Information 3 - dMIQE Checklist

Supporting Information 4 - Sampling Loss Model

Supporting Information 5 - Shedding Rates Summary

Supporting Information 6 - Decay Rates Summary

## Acknowledgement

We thank Mariana Jimenez, Celine de Jong, Vince Christian, Rosie Arathoon, Erika Camacho, Aaron Rosenfeld and Cailley Gerald-Yamasaki for assistance with experimental set-up and sample collection. We are also grateful to Vince and Beth Christian for field work accommodations. Thank you to Cole Citrenbaum of the Stanford Statistical Consulting Services for guidance on model selection for shedding rate comparisons, to Sarah Lowry for assistance with photochemistry calculations, to Vincent DiPietri for helping collect absorbance measurements, and to Royal Kopperud and Carine Sauquet for laboratory and administrative support. This is NOAA’s Pacific Marine Environmental Laboratory (PMEL) contribution number 5912. Zachary Gold was supported by NOAA’s PMEL and the NOAA’s Omics Program.

## Notes

### Competing Interest Statement

The authors have declared no competing interest.

## References

(1) Edwards, M. S. Kelp Forests Under Threat and the Loss of Ecosystem Services They Provide. Annu. Rev. Ecol. Evol. Syst. 2025, 56 (1), 365–392. 10.1146/annurev-ecolsys-102723-055806

(2) Connell, S.; Russell, B.; Turner, D.; Shepherd, S.; Kildea, T.; Miller, D.; Airoldi, L.; Cheshire, A. Recovering a Lost Baseline: Missing Kelp Forests from a Metropolitan Coast. Mar. Ecol. Prog. Ser. 2008, 360, 63–72. 10.3354/meps07526

(3) Fowles, A. E.; Edgar, G. J.; Hill, N.; Stuart-Smith, R. D.; Kirkpatrick, J. B. An Experimental Assessment of Impacts of Pollution Sources on Sessile Biota in a Temperate Urbanised Estuary. Mar. Pollut. Bull. 2018, 133, 209–217. 10.1016/j.marpolbul.2018.05.040

(4) Fowles, A. E.; Stuart-Smith, R. D.; Stuart-Smith, J. F.; Hill, N. A.; Kirkpatrick, J. B.; Edgar, G. J. Effects of Urbanisation on Macroalgae and Sessile Invertebrates in Southeast Australian Estuaries. Est. Coast. Shelf Sci. 2018, 205, 30–39. 10.1016/j.ecss.2018.02.010

(5) Oyarzo-Miranda, C.; Latorre, N.; Meynard, A.; Rivas, J.; Bulboa, C.; Contreras-Porcia, L. Coastal Pollution from the Industrial Park Quintero Bay of Central Chile: Effects on Abundance, Morphology, and Development of the Kelp Lessonia Spicata (Phaeophyceae). PLOS ONE 2020, 15 (10), e0240581. 10.1371/journal.pone.0240581

(6) Smale, D. A. Impacts of Ocean Warming on Kelp Forest Ecosystems. New Phytol. 2020, 225 (4), 1447–1454. 10.1111/nph.16107

(7) Krumhansl, K. A.; Okamoto, D. K.; Rassweiler, A.; Novak, M.; Bolton, J. J.; Cavanaugh, K. C.; Connell, S. D.; Johnson, C. R.; Konar, B.; Ling, S. D.; Micheli, F.; Norderhaug, K. M.; Pérez-Matus, A.; Sousa-Pinto, I.; Reed, D. C.; Salomon, A. K.; Shears, N. T.; Wernberg, T.; Anderson, R. J.; Barrett, N. S.; Buschmann, A. H.; Carr, M. H.; Caselle, J. E.; Derrien-Courtel, S.; Edgar, G. J.; Edwards, M.; Estes, J. A.; Goodwin, C.; Kenner, M. C.; Kushner, D. J.; Moy, F. E.; Nunn, J.; Steneck, R. S.; Vásquez, J.; Watson, J.; Witman, J. D.; Byrnes, J. E. K. Global Patterns of Kelp Forest Change over the Past Half-Century. Proc. Natl. Acad. Sci. U.S.A. 2016, 113 (48), 13785–13790. 10.1073/pnas.1606102113

(8) Wernberg, T.; Krumhansl, K.; Filbee-Dexter, K.; Pedersen, M. F. Status and Trends for the World’s Kelp Forests. In World Seas: An Environmental Evaluation; Academic Press, 2019; pp 57–78. 10.1016/B978-0-12-805052-1.00003-6

(9) Galloway, A. W. E.; Gravem, S. A.; Kobelt, J. N.; Heady, W. N.; Okamoto, D. K.; Sivitilli, D. M.; Saccomanno, V. R.; Hodin, J.; Whippo, R. Sunflower Sea Star Predation on Urchins Can Facilitate Kelp Forest Recovery. *Proc. R. Soc.*, B 2023, 290 (1993), 20221897. 10.1098/rspb.2022.1897

(10) Prentice, M. B.; Crandall, G. A.; Chan, A. M.; Davis, K. M.; Hershberger, P. K.; Finke, J. F.; Hodin, J.; McCracken, A.; Kellogg, C. T. E.; Clemente-Carvalho, R. B. G.; Prentice, C.; Zhong, K. X.; Harvell, C. D.; Suttle, C. A.; Gehman, A.-L. M. Vibrio Pectenicida Strain FHCF-3 Is a Causative Agent of Sea Star Wasting Disease. *Nat*. Ecol. Evol. 2025, 9 (9), 1739–1751. 10.1038/s41559-025-02797-2

(11) Gravem, S. A.; Heady, W. N.; Saccomanno, V. R.; Alvstad, K. F.; Gehman, A. L. M.; Freirson, T.; Hamilton, S. L. Pycnopodia Helianthoides (Amended Version of 2020 Assessment). The IUCN Red List of Threatened Species 2021: e.T178290276A197818455. 2021. 10.2305/IUCN.UK.2021-1.RLTS.T178290276A197818455.en

(12) Hewson, I.; Button, J. B.; Gudenkauf, B. M.; Miner, B.; Newton, A. L.; Gaydos, J. K.; Wynne, J.; Groves, C. L.; Hendler, G.; Murray, M.; Fradkin, S.; Breitbart, M.; Fahsbender, E.; Lafferty, K. D.; Kilpatrick, A. M.; Miner, C. M.; Raimondi, P.; Lahner, L.; Friedman, C. S.; Daniels, S.; Haulena, M.; Marliave, J.; Burge, C. A.; Eisenlord, M. E.; Harvell, C. D. Densovirus Associated with Sea-Star Wasting Disease and Mass Mortality. Proc. Natl. Acad. Sci. U.S.A. 2014, 111 (48), 17278–17283. 10.1073/pnas.1416625111

(13) Dawson, M. N.; Duffin, P. J.; Giakoumis, M.; Schiebelhut, L. M.; Beas-Luna, R.; Bosley, K. L.; Castilho, R.; Ewers-Saucedo, C.; Gavenus, K. A.; Keller, A.; Konar, B.; Largier, J. L.; Lorda, J.; Miner, C. M.; Moritsch, M. M.; Navarrete, S. A.; Traiger, S. B.; Turner, M. S.; Wares, J. P. A Decade of Death and Other Dynamics: Deepening Perspectives on the Diversity and Distribution of Sea Stars and Wasting. Biol. Bull. 2023, 244 (3), 143–163. 10.1086/727969

(14) Hodin, J.; Anteau, F. P.; Ashcraft, B. F.; Brito, M.; Curliss, F.; Kalytiak-Davis, A. R.; Peng, J. S.; Schwab, C. J.; Valdez, V. V.; Weertman, W. Star Power: Early Life Stages of an Endangered Sea Star Are Robust to Current and near-Future Warming. PLOS ONE 2025, 20 (9), e0318879. 10.1371/journal.pone.0318879

(15) Mancuso, R. T.; Gravem, S. A.; Campbell, R. S.; Hunter, N.; Raimondi, P.; Galloway, A. W. E.; Kroeker, K. J. Sunflower Sea Star Chemical Cues Locally Reduce Kelp Consumption by Eliciting a Flee Response in Red Sea Urchins. *Proc. R. Soc.*, B 2025, 292 (2050), 20250949. 10.1098/rspb.2025.0949

(16) Lowry, D.; Wright, S.; Neuman, M.; Stevenson, D.; Hyde, J.; Lindeberg, M.; Tolimieri, N.; Lonhart, S.; Traiger, S.; Gustafson, R. Endangered Species Act Status Review Report: Sunflower Sea Star (Pycnopodia Helianthoides); Final Report to the National Marine Fisheries Service, Office of Protected Resources.; 2022.

(17) Taberlet, P.; Bonin, A.; Zinger, L.; Coissac, E. Environmental DNA: For Biodiversity Research and Monitoring, 1st ed.; Oxford University Press: Oxford, United Kingdom, 2018.

(18) Stewart, K. A. Understanding the Effects of Biotic and Abiotic Factors on Sources of Aquatic Environmental DNA. Biodivers Conserv 2019, 28 (5), 983–1001. 10.1007/s10531-019-01709-8

(19) Miya, M. Environmental DNA Metabarcoding: A Novel Method for Biodiversity Monitoring of Marine Fish Communities. Annual Review of Marine Science 2022, 14 (1), 161–185. 10.1146/annurev-marine-041421-082251

(20) Williams, M. A.; Bracken, F. S. A.; Idelegbagbon, O.; Parle-McDermott, A. Environmental DNA as a Tool for Single Species Detection. In Biosensors for the Marine Environment: Present and Future Challenges; Regan, F., Hansen, P.-D., Barceló, D., Eds.; Springer International Publishing: Cham, 2023; pp 63–94. 10.1007/698_2022_956

(21) Iacaruso, N. J.; Reves, O. P.; Merkelz, S. J.; Waldrep, C. L.; Davis, M. A. A Systematic Review Evaluating the Performance of eDNA Methods Relative to Conventional Methods for Biodiversity Monitoring. Ecography 2026, *n/a* (n/a), e07952. 10.1002/ecog.07952

(22) Gold, Z.; Robinson, K. M.; Gehman, A.-L. M.; Shea, M. M.; Lemay, M. A.; Weinrich, J.; Kellogg, C. T. E.; Clemente-Carvalho, R. B. G.; Schiebelhut, L. M.; Boehm, A. B.; Kidd, A.; Kim, A.; Hodin, J.; Dawson, M.; McAllister, S. M. Molecular Star Gazing: Development and Validation of an Environmental DNA Assay for the Imperiled Sunflower Sea Star (Pycnopodia Helianthoides). bioRxiv May 12, 2026, p 2026.05.07.723600. 10.64898/2026.05.07.723600

(23) Harrison, J. B.; Sunday, J. M.; Rogers, S. M. Predicting the Fate of eDNA in the Environment and Implications for Studying Biodiversity. *Proc. R. Soc.*, B 2019, 286, 20191409. 10.1098/rspb.2019.1409

(24) Scriver, M.; Zaiko, A.; Pochon, X.; von Ammon, U. Harnessing Decay Rates for Coastal Marine Biosecurity Applications: A Review of Environmental DNA and RNA Fate. Environmental DNA 2023, 5 (5), 960–972. 10.1002/edn3.405

(25) Andruszkiewicz Allan, E.; Zhang, W. G.; C. Lavery, A.; F. Govindarajan, A. Environmental DNA Shedding and Decay Rates from Diverse Animal Forms and Thermal Regimes. Environmental DNA 2021, 3, 492–514. 10.1002/edn3.141

(26) Collins, R. A.; Wangensteen, O. S.; O’Gorman, E. J.; Mariani, S.; Sims, D. W.; Genner, M. J. Persistence of Environmental DNA in Marine Systems. COMMUNICATIONS BIOLOGY 2018, 1. 10.1038/s42003-018-0192-6

(27) Lancaster, E. R.; Grey, E. K.; Brady, D. C.; Frederich, M. Squishy and Crunchy Invasive Invertebrates: Environmental DNA Is Not Shed Equally. Environmental DNA 2025, 7 (2). 10.1002/edn3.70072

(28) Jo, T.; Murakami, H.; Masuda, R.; Sakata, M. K.; Yamamoto, S.; Minamoto, T. Rapid Degradation of Longer DNA Fragments Enables the Improved Estimation of Distribution and Biomass Using Environmental DNA. Mol Ecol Resour 2017, 17 (6), e25–e33. 10.1111/1755-0998.12685

(29) Ballmer, E.; McNeill, K.; Deiner, K. Potential Role of Photochemistry in Environmental DNA Degradation. Environ. Sci. Technol. Lett. 2024, 11 (12), 1284–1295. 10.1021/acs.estlett.4c00704

(30) Andruszkiewicz, E. A.; Sassoubre, L. M.; Boehm, A. B. Persistence of Marine Fish Environmental DNA and the Influence of Sunlight. PLOS ONE 2017, 12 (9), e0185043. 10.1371/journal.pone.0185043

(31) Sigsgaard, E. E.; Nielsen, I. B.; Bach, S. S.; Lorenzen, E. D.; Robinson, D. P.; Knudsen, S. W.; Pedersen, M. W.; Al Jaidah, M.; Orlando, L.; Willerslev, E.; Moller, P. R.; Thomsen, P. F. Population Characteristics of a Large Whale Shark Aggregation Inferred from Seawater Environmental DNA. NATURE ECOLOGY & EVOLUTION 2017, 1 (1). 10.1038/s41559-016-0004

(32) Borchardt, M. A.; Boehm, A. B.; Salit, M.; Spencer, S. K.; Wigginton, K. R.; Noble, R. T. The Environmental Microbiology Minimum Information (EMMI) Guidelines: qPCR and dPCR Quality and Reporting for Environmental Microbiology. Environ. Sci. Technol. 2021, 55 (15), 10210–10223. 10.1021/acs.est.1c01767

(33) The dMIQE Group; Huggett, J. F. The Digital MIQE Guidelines Update: Minimum Information for Publication of Quantitative Digital PCR Experiments for 2020. Clin Chem 2020, 66 (8), 1012–1029. 10.1093/clinchem/hvaa125

(34) Shea, M. M.; Boehm, A. B. Meghanmshea/Stanford-Boehm_Pycno_v1_Collection_Protocol_Sterivex_GravityBag: V0.0.0. 2026. 10.5281/zenodo.22834297

(35) Shea, M. M.; Boehm, A. B. DNA Extraction Protocol from Sterivex Filters. 2023. 10.17504/protocols.io.ewov1qyyygr2/v1

(36) Shea, M. M.; Boehm, A. B. Meghanmshea/Stanford-Boehm_Pycno_v1_Extraction_Protocol_BeBOP: V1.0.0. 2026. 10.5281/zenodo.22663570

(37) Shea, M. M.; Kubiatko, L.; Boehm, A. B. Meghanmshea/Stanford-Boehm_Pycno_v1_ddPCR_Protocol_BeBOP: V1.0.0, 2026. 10.5281/zenodo.20057581

(38) Shea, M. M.; Boehm, A. B. Coastal Environmental DNA Sampling & Gravity Filtration Protocol. 2023. 10.17504/protocols.io.bp2l69y7klqe/v2

(39) Spens, J.; Evans, A. R.; Halfmaerten, D.; Knudsen, S. W.; Sengupta, M. E.; Mak, S. S. T.; Sigsgaard, E. E.; Hellstrom, M. Comparison of Capture and Storage Methods for Aqueous Macrobial eDNA Using an Optimized Extraction Protocol: Advantage of Enclosed Filter. Methods in Ecology and Evolution 2017, 8 (5), 635–645. 10.1111/2041-210X.12683

(40) Samuel, R. M.; Meyer, R.; Buttigieg, P. L.; Davies, N.; Jeffery, N. W.; Meyer, C.; Pavloudi, C.; Pitz, K. J.; Sweetlove, M.; Theroux, S.; van de Kamp, J.; Watts, A. Toward a Global Public Repository of Community Protocols to Encourage Best Practices in Biomolecular Ocean Observing and Research. Front. Mar. Sci. 2021, 8. 10.3389/fmars.2021.758694

(41) Cao, Y.; Griffith, J. f.; Dorevitch, S.; Weisberg, S. b. Effectiveness of qPCR Permutations, Internal Controls and Dilution as Means for Minimizing the Impact of Inhibition While Measuring Enterococcus in Environmental Waters. Journal of Applied Microbiology 2012, 113 (1), 66–75. 10.1111/j.1365-2672.2012.05305.x

(42) Sassoubre, L. M.; Yamahara, K. M.; Gardner, L. D.; Block, B. A.; Boehm, A. B. Quantification of Environmental DNA (eDNA) Shedding and Decay Rates for Three Marine Fish. Environ. Sci. Technol. 2016, 50 (19), 10456–10464. 10.1021/acs.est.6b03114

(43) Sansom, B. J.; Sassoubre, L. M. Environmental DNA (eDNA) Shedding and Decay Rates to Model Freshwater Mussel eDNA Transport in a River. ENVIRONMENTAL SCIENCE & TECHNOLOGY 2017, 51 (24), 14244–14253. 10.1021/acs.est.7b05199

(44) Bates, D.; Mächler, M.; Bolker, B.; Walker, S. Fitting Linear Mixed-Effects Models Using **Lme4**. J. Stat. Soft. 2015, 67 (1). 10.18637/jss.v067.i01

(45) Kuznetsova, A.; Brockhoff, P. B.; Christensen, R. H. B. **lmerTest** Package: Tests in Linear Mixed Effects Models. J. Stat. Soft. 2017, 82 (13). 10.18637/jss.v082.i13

(46) Grant, M. J.; Booth, A. A Typology of Reviews: An Analysis of 14 Review Types and Associated Methodologies. Health Information & Libraries Journal 2009, 26 (2), 91–108. 10.1111/j.1471-1842.2009.00848.x

(47) Jo, T.; Minamoto, T. Complex Interactions between Environmental DNA (eDNA) State and Water Chemistries on eDNA Persistence Suggested by Meta-Analyses. Molecular Ecology Resources 2021, 21 (5), 1490–1503. 10.1111/1755-0998.13354

(48) Lamb, P. D.; Fonseca, V. G.; Maxwell, D. L.; Nnanatu, C. C. Systematic Review and Meta-Analysis: Water Type and Temperature Affect Environmental DNA Decay. Molecular Ecology Resources 2022, 22 (7), 2494–2505. 10.1111/1755-0998.13627

(49) Waters, T.; Langlois, K.; Gold, Z.; Theroux, S.; Eagle, R. A. Hidden in Plain Sight: The Invasive Macroalga Caulerpa Prolifera Evades Detection by Environmental DNA Methods. Environmental DNA 2024, 6 (1). 10.1002/edn3.496

(50) Klymus, K. E.; Ramos, D. V. R.; Thompson, N. L.; Richter, C. A. Development and Testing of Species-Specific Quantitative PCR Assays for Environmental DNA Applications. Journal of Visualized Experiments (JoVE) 2020, No. 165, e61825. 10.3791/61825

(51) Brasseale, E.; Adams, N.; Allan, E. A.; Jacobson, E. K.; Kelly, R. P.; Liu, O. R.; Moore, S.; Shaffer, M.; Xiong, J.; Parsons, K. Marine eDNA Production and Loss Mechanisms. *Journal of Geophysical Research: Oceans* 2025, 130 (4). 10.1029/2024JC021643

(52) Chung, A.; Kam, Y. C.; Schunter, C. Novel Assay for Endangered Hong Kong Grouper (Epinephelus Akaara) to Assess eDNA Shedding, Decay, and Population Status. Environmental DNA 2026, 8 (2). 10.1002/edn3.70264

(53) Cowart, D. A.; Murphy, K. R.; Cheng, C.-H. C. Metagenomic Sequencing of Environmental DNA Reveals Marine Faunal Assemblages from the West Antarctic Peninsula. MARINE GENOMICS 2018, 37, 148–160. 10.1016/j.margen.2017.11.003

(54) Dass, M.; Sherman, C.; van Oorschot, R.; Hartman, D.; Carter, G.; Durdle, A. A Preliminary Study on Detecting Human DNA in Aquatic Environments: Potential of eDNA in Forensics. FORENSIC SCIENCE INTERNATIONAL-GENETICS 2025, 74. 10.1016/j.fsigen.2024.103155

(55) Ellis, M. R.; Clark, Z. S. R.; Treml, E. A.; Brown, M. S.; Matthews, T. G.; Pocklington, J. B.; Stafford-Bell, R. E.; Bott, N. J.; Nai, Y. H.; Miller, A. D.; Sherman, C. D. H. Detecting Marine Pests Using Environmental DNA and Biophysical Models. Science of the Total Environment 2022, 816. 10.1016/j.scitotenv.2021.151666

(56) Forsstrom, T.; Vasemagi, A. Can Environmental DNA (eDNA) Be Used for Detection and Monitoring of Introduced Crab Species in the Baltic Sea? MARINE POLLUTION BULLETIN 2016, 109 (1), 350–355. 10.1016/j.marpolbul.2016.05.054

(57) Holman, L. E.; Chng, Y.; Rius, M. How Does eDNA Decay Affect Metabarcoding Experiments? Environmental DNA 2022, 4 (1), 108–116. 10.1002/edn3.201

(58) Jo, T.; Murakami, H.; Yamamoto, S.; Masuda, R.; Minamoto, T. Effect of Water Temperature and Fish Biomass on Environmental DNA Shedding, Degradation, and Size Distribution. ECOLOGY AND EVOLUTION 2019, 9 (3), 1135–1146. 10.1002/ece3.4802

(59) Jo, T.; Arimoto, M.; Murakami, H.; Masuda, R.; Minamoto, T. Estimating Shedding and Decay Rates of Environmental Nuclear DNA with Relation to Water Temperature and Biomass. Environmental DNA 2020, 2 (2), 140–151. 10.1002/edn3.51

(60) Jo, T.; Tsuri, K.; Hirohara, T.; Yamanaka, H. Warm Temperature and Alkaline Conditions Accelerate Environmental RNA Degradation. Environmental DNA 2023, 5 (5), 836–848. 10.1002/edn3.334

(61) Kirtane, A.; Wieczorek, D.; Noji, T.; Baskin, L.; Ober, C.; Plosica, R.; Chenoweth, A.; Lynch, K.; Sassoubre, L. Quantification of Environmental DNA (eDNA) Shedding and Decay Rates for Three Commercially Harvested Fish Species and Comparison between eDNA Detection and Trawl Catches. Environmental DNA 2021, 3 (6), 1142–1155. 10.1002/edn3.236

(62) Kutti, T.; Johnsen, I. A.; Skaar, K. S.; Ray, J. L.; Husa, V.; Dahlgren, T. G. Quantification of eDNA to Map the Distribution of Cold-Water Coral Reefs. FRONTIERS IN MARINE SCIENCE 2020, 7. 10.3389/fmars.2020.00446

(63) Kwong, S. L. T.; Villacorta-Rath, C.; Doyle, J.; Uthicke, S. Quantifying Shedding and Degradation Rates of Environmental DNA (eDNA) from Pacific Crown-of-Thorns Seastar (Acanthaster Cf. Solaris). Marine Biology 2021, 168 (6). 10.1007/s00227-021-03896-x

(64) Lopez, M. L. D.; Rolheiser, K. C.; Etzkorn, J.; Imbery, J. J.; Lemay, M. A.; Gimenez, I.; Helbing, C. C. Impact of CO2-Induced Aquatic Acidification on Environmental DNA and RNA Shedding and Persistence. Environmental DNA 2025, 7 (4). 10.1002/edn3.70158

(65) McCartin, L. J.; Vohsen, S. A.; Ambrose, S. W.; Layden, M.; McFadden, C. S.; Cordes, E. E.; McDermott, J. M.; Herrera, S. Temperature Controls eDNA Persistence across Physicochemical Conditions in Seawater. Environmental Science and Technology 2022, 56 (12), 8629–8639. 10.1021/acs.est.2c01672

(66) Minamoto, T.; Fukuda, M.; Katsuhara, K. R.; Fujiwara, A.; Hidaka, S.; Yamamoto, S.; Takahashi, K.; Masuda, R. Environmental DNA Reflects Spatial and Temporal Jellyfish Distribution. PLoS ONE 2017, 12 (2), e0173073. 10.1371/journal.pone.0173073

(67) Morrissey, S. J.; Jerry, D. R.; Kingsford, M. J. Genetic Detection and a Method to Study the Ecology of Deadly Cubozoan Jellyfish. Diversity 2022, 14 (12). 10.3390/d14121139

(68) Qian, T.; Shan, X.; Wang, W.; Jin, X. Effects of Temperature on the Timeliness of eDNA/eRNA: A Case Study of Fenneropenaeus Chinensis. Water (Switzerland) 2022, 14 (7). 10.3390/w14071155

(69) Saito, T.; Doi, H. Degradation Modeling of Water Environmental DNA: Experiments on Multiple DNA Sources in Pond and Seawater. Environmental DNA 2021, 3 (4), 850–860. 10.1002/edn3.192

(70) Skinner, E.; Hamann, M.; Strugnell, J.; Rath, C. Investigating the Limits of Detection of Environmental DNA From Loggerhead and Hawksbill Turtles. Aquatic Conservation: Marine and Freshwater Ecosystems 2025, 35 (8). 10.1002/aqc.70199

(71) Thomsen, P. F.; Kielgast, J.; Iversen, L. L.; Møller, P. R.; Rasmussen, M.; Willerslev, E. Detection of a Diverse Marine Fish Fauna Using Environmental DNA from Seawater Samples. PLOS ONE 2012, 7 (8), e41732. 10.1371/journal.pone.0041732

(72) Weltz, K.; Lyle, J. M.; Ovenden, J.; Morgan, J. A. T.; Moreno, D. A.; Semmens, J. M. Application of Environmental DNA to Detect an Endangered Marine Skate Species in the Wild. PLOS ONE 2017, 12 (6). 10.1371/journal.pone.0178124

(73) Wood, S. A.; Biessy, L.; Latchford, J. L.; Zaiko, A.; von Ammon, U.; Audrezet, F.; Cristescu, M. E.; Pochon, X. Release and Degradation of Environmental DNA and RNA in a Marine System. SCIENCE OF THE TOTAL ENVIRONMENT 2020, 704. 10.1016/j.scitotenv.2019.135314

(74) Beattie, R. E.; Helbing, C. C.; Imbery, J. J.; Klymus, K. E.; Lopez Duran, J.; Richter, C. A.; Thambirajah, A. A.; Thompson, N.; Edwards, T. M. A Nitrifier-Enriched Microbial Community Contributes to the Degradation of Environmental DNA. Environmental DNA 2023, 5 (6), 1473–1483. 10.1002/edn3.469

(75) Ibáñez de Aldecoa, A. L.; Zafra, O.; González-Pastor, J. E. Mechanisms and Regulation of Extracellular DNA Release and Its Biological Roles in Microbial Communities. Front. Microbiol. 2017, 8. 10.3389/fmicb.2017.01390

(76) Mauvisseau, Q.; Harper, L. R.; Sander, M.; Hanner, R. H.; Kleyer, H.; Deiner, K. The Multiple States of Environmental DNA and What Is Known about Their Persistence in Aquatic Environments. Environ. Sci. Technol. 2022, 56 (9), 5322–5333. 10.1021/acs.est.1c07638

(77) Mishonov, A. V.; Boyer, T. P.; Baranova, O. K.; Bouchard, C. N.; Cross, S. L.; Garcia, H. E.; Locarnini, R. A.; Paver, C. R.; Reagan, J. R.; Wang, Z.; Seidov, D.; Grodsky, A. I.; Beauchamp, J. G. World Ocean Database 2023. NOAA atlas NESDIS ; 97 2024. 10.25923/Z885-H264

(78) Hamel, J.-F.; Jobson, S.; Caulier, G.; Mercier, A. Evidence of Anticipatory Immune and Hormonal Responses to Predation Risk in an Echinoderm. Sci Rep 2021, 11 (1), 10691. 10.1038/s41598-021-89805-0

(79) Thimmappa, R.; Wang, S.; Zheng, M.; Misra, R. C.; Huang, A. C.; Saalbach, G.; Chang, Y.; Zhou, Z.; Hinman, V.; Bao, Z.; Osbourn, A. Biosynthesis of Saponin Defensive Compounds in Sea Cucumbers. Nat Chem Biol 2022, 18 (7), 774–781. 10.1038/s41589-022-01054-y

(80) Maruyama, A.; Nakamura, K.; Yamanaka, H.; Kondoh, M.; Minamoto, T. The Release Rate of Environmental DNA from Juvenile and Adult Fish. PLoS One 2014, 9 (12), e114639. 10.1371/journal.pone.0114639

(81) López-Nandam, E. H.; Story, L. T.; Evin, M.; Witherly, J.; Upton, M.; Krol, L.; Dunker, F.; Lev, K. Growth, Survival, and Fitness in the First Year of Life for Pycnopodia Helianthoides under Different Larval Densities. bioRxiv April 6, 2026, p 2026.04.02.716152. 10.64898/2026.04.02.716152

(82) Andruszkiewicz, E. A.; Koseff, J. R.; Fringer, O. B.; Ouellette, N. T.; Lowe, A. B.; Edwards, C. A.; Boehm, A. B. Modeling Environmental DNA Transport in the Coastal Ocean Using Lagrangian Particle Tracking. Front. Mar. Sci. 2019, 6, 477. 10.3389/fmars.2019.00477

(83) Uthicke, S.; Robson, B.; Doyle, J. R.; Logan, M.; Pratchett, M. S.; Lamare, M. Developing an Effective Marine eDNA Monitoring: eDNA Detection at Pre-Outbreak Densities of Corallivorous Seastar (*Acanthaster* Cf. Solaris). Science of The Total Environment 2022, 851, 158143. 10.1016/j.scitotenv.2022.158143

