## Supporting Information 1 - Additional Methodological Details for "Sunflower sea star (*Pycnopodia helianthoides*) environmental DNA (eDNA) shedding and decay: Influence of sunlight and comparison to other marine organisms"

### *1. Additional Information for dMIQE Reporting Guidelines*

Bio-Rad ddPCR Supermix for probes, no dUTP (catalog number 186-3024, Bio-Rad, Hercules, CA, USA) came from lots 64705079 and 64707759.

Our experimental shedding samples had a range of 19,159 to 40,146 partitions (average: 37,665 ± 2427) and copies per partition (λ) from 0 to 0.882 (average: 0.108 ± 0.157). Our experimental decay samples had a range of 10,711 to 50,284 partitions (average: 34,204 ± 5953) and copies per partition from 0.048 to 2.070 (average: 0.628 ± 0.385). The reported partition volume for the QX200 Droplet Digital PCR System (Bio-Rad, Hercules, CA, USA) is 0.00085 µL.

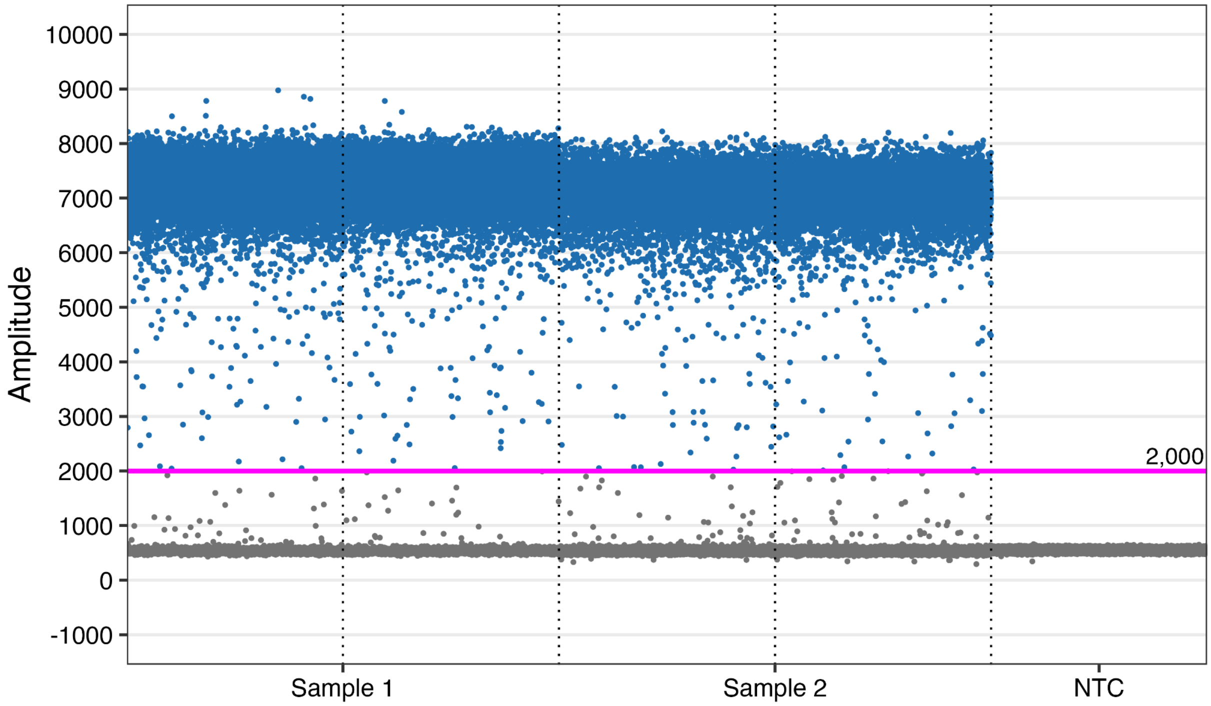

**Figure S1:** Example fluorescence plot for samples (first four columns) and negative controls (last column). The threshold used for the given plate is indicated in red.

*2. Time to Steady State Derivation*

In order to estimate when steady state might be reached in this system, we solved the full shedding equation for when C = $C_{ss}$*0.99, where V is the volume of the aquaria (L), C is the concentration of eDNA (copies/L), t is the time elapsed (hr), S is the eDNA shedding rate (copies/hr), k is the decay rate constant ($hr^{-1}$), $C_{ss}$ is the concentration of eDNA at steady state (copies/L), and $t_{ss}$ is the time to steady state (hr):

1. Time rate of change of concentration of eDNA equation: $C(t) = (C_0 - \frac{S}{Vk}) e^{-kt} + \frac{S}{Vk}$
2. To estimate steady state, we want to know when the concentration approaches the concentration at steady state, or when $C = 0.99 \cdot C_{ss}$.
3. From Equation 4, we know that $C_{ss} = \frac{S}{Vk}$, so we want to solve for $C = 0.99 \frac{S}{Vk}$
4. $0.99 \frac{S}{Vk} = (C_0 - \frac{S}{Vk}) e^{-kt_{ss}} + \frac{S}{Vk}$
5. Assuming $C_0 = 0$, $0.99 \frac{S}{Vk} = - \frac{S}{Vk} e^{-kt_{ss}} + \frac{S}{Vk}$
6. $0.99 = - e^{-kt_{ss}} + 1$
7. $e^{-kt_{ss}} = (1 - 0.99)$
8. $- kt_{ss} = ln(0.01)$
9. $t_{ss} = \frac{-ln(0.01)}{k}$

*3. Calculation of Daily Average UVA and UVB Irradiance*

To calculate daily average UVA and UVB irradiance, we followed the approach from Maraccini et al. [1], but we used measured irradiance rather than the Simple Model of the Atmospheric Radiative Transfer of Sunshine (SMARTS). For each radiometer measurement collected over the course of the study, irradiance of wavelengths from 280 - 315 nm was summed to obtain the UVB irradiance and irradiance of wavelengths from 316 - 400 nm was summed to obtain the UVA irradiance (W/m$^2$). Measurements were collected from sunrise to sunset; to calculate average UVA and UVB irradiance on a given day, we also needed to account for conditions between sunset and sunrise, when we assumed irradiance was 0. To do so, we extended the sampling interval each day before and after sunrise, and set irradiance to 0 at those time points (see Figures S1). Then, we calculated the average UVA and UVB irradiance across all original measurements and extended time points for a given day. This gave us the average UVA and UVB irradiance for each day of our experiment, for our treatment and control conditions (Figure S2, Table S1). Finally, we calculated the average UVA and UVB irradiance across the four full days of our experiment by taking the mean of the daily averages.

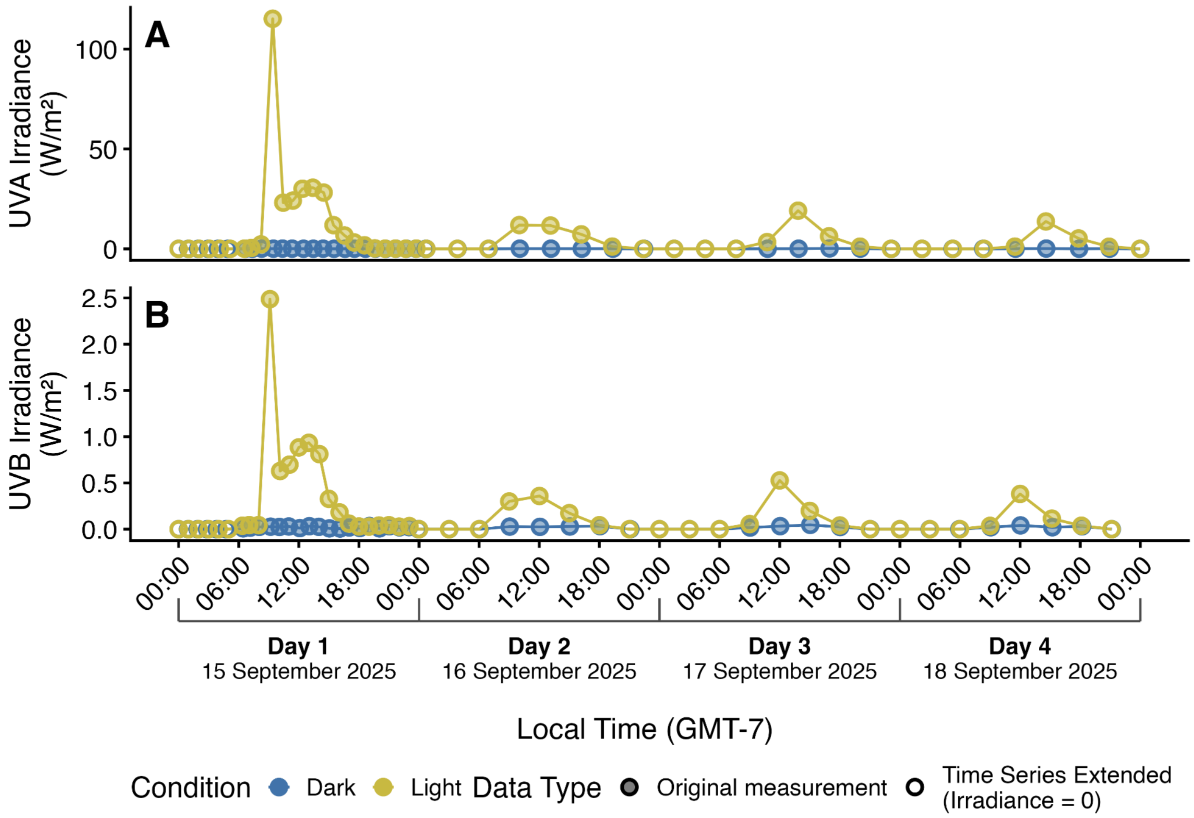

**Figure S2:** UVA (A) and UVB (B) irradiance measured using the radiometer at the surface of each experimental condition (light and dark) as a function of local time; filled points are directly measured values and open points are where we have extended our time series to account for unsampled night-time hours (irradiance = 0), which are used for our daily averages.

Then, we adjusted these measurements to account for the attenuation of UVA and UVB into the enteral feeding pouches. We derived an  attenuation factor by comparing the radiometer measurements of incident sunlight irradiance and sunlight irradiance after it passed through the pouch material (Figure S3), using the following equation (Equation S1):

$$A = 1 + \frac{E_{sunlight + pouch, \lambda} - E_{sunlight, \lambda}}{E_{sunlight, \lambda}}$$

where $A$ is the attenuation factor and $E_{x, \lambda}$ is the measured irradiance of a given wavelength ($W/m^2$) for a given condition.

To calculate the UVA and UVB incident on the surface of water inside the pouch, we multiplied measured irradiance at each wavelength by its corresponding attenuation factor ("Passing Through Pouch" in Table S1). We calculated the average percent attenuation across the UVA and UVB spectra with the following equation (Equation S2) :

$$Percent\ Attenuation = \frac{\sum_{\lambda_1}^{\lambda_2} E_{sunlight + pouch, \lambda} - \sum_{\lambda_1}^{\lambda_2} E_{sunlight, \lambda}}{\sum_{\lambda_1}^{\lambda_2} E_{sunlight, \lambda}}$$

where $\lambda_1$ and $\lambda_2$ are the lower and upper bounds of the spectra of interest. We found an attenuation across the wavelengths that comprise each spectra of 14% UVA and 16.7% UVB.

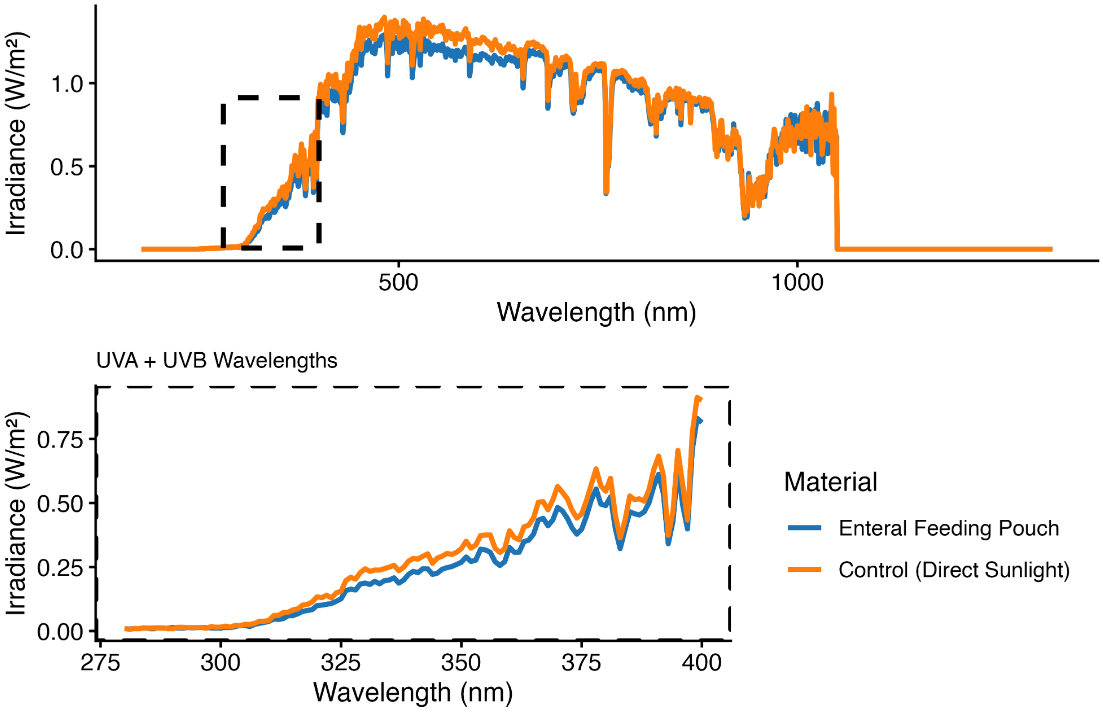

**Figure S3:** Measured irradiance values of incident sunlight and sunlight passing through the enteral feeding pouch, as used to calculate the attenuation factor per wavelength

Following Anderson & Boehm [2], we calculated the depth-averaged light intensity accounting for light screening by the experimental solution within the pouch using the following equation (Equation S3):

$$\langle E_{0,\lambda}\rangle_z = \frac{1-10^{-\alpha_\lambda z}}{2.303\times\alpha_\lambda z}E_{0,\lambda}(0)$$

where $\langle E_{0,\lambda}\rangle_z$ is the average spectral irradiance (W/m$^2$) over a given depth z (m), $E_{0,\lambda}(0)$ is the irradiance at depth 0 (just below the water surface), and $\alpha_\lambda$ is the decadic absorption coefficient (m$^{-1}$) calculated by dividing the absorbance measured by the radiometer (Figure S4) by the cuvette width (0.01 m). Depth (0.058 m) was estimated by measuring the enteral feeding pouches when full and floating in water, although we note that they are not uniform shapes nor were they positioned perfectly perpendicular in the water bath. Light absorption of the water from the Sunflower Star Laboratory flow-through system was measured  using a Cary 60 UV-Vis spectrophotometer (Agilent Technologies, Santa Clara, CA) . Depth-averaged values are summarized in Table S1. An example of the relative impact of the pouch material attenuation and water absorption on irradiance is shown in Figure S5.

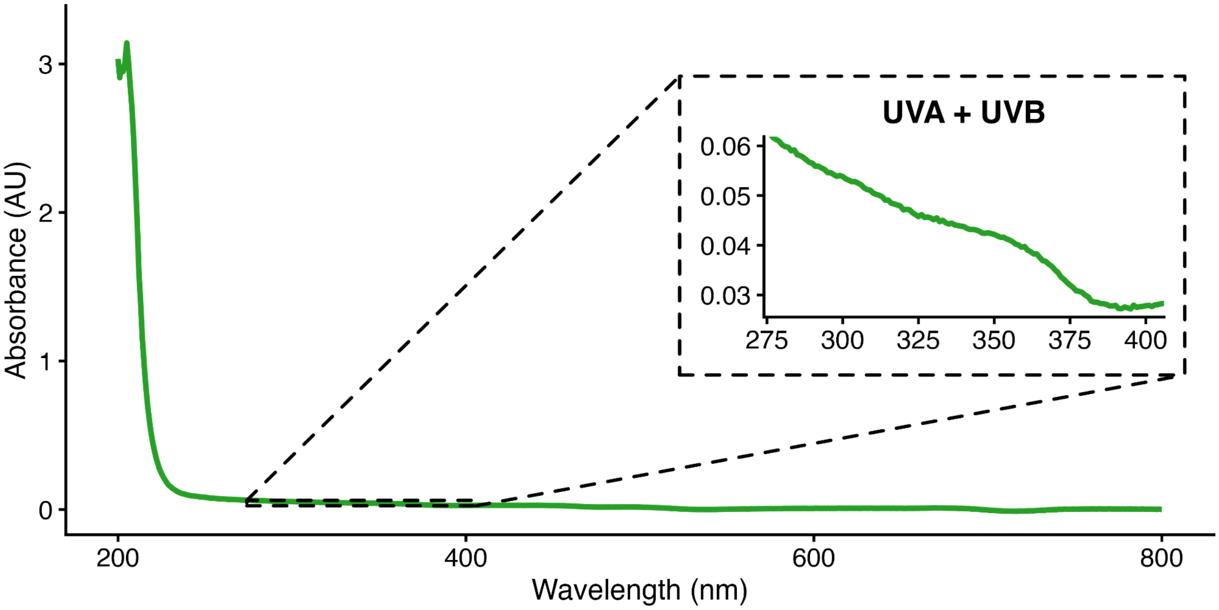

**Figure S4:** Measured absorbance values of experimental water, as used to calculate the depth-averaged light intensity accounting for light screening by the experimental solution

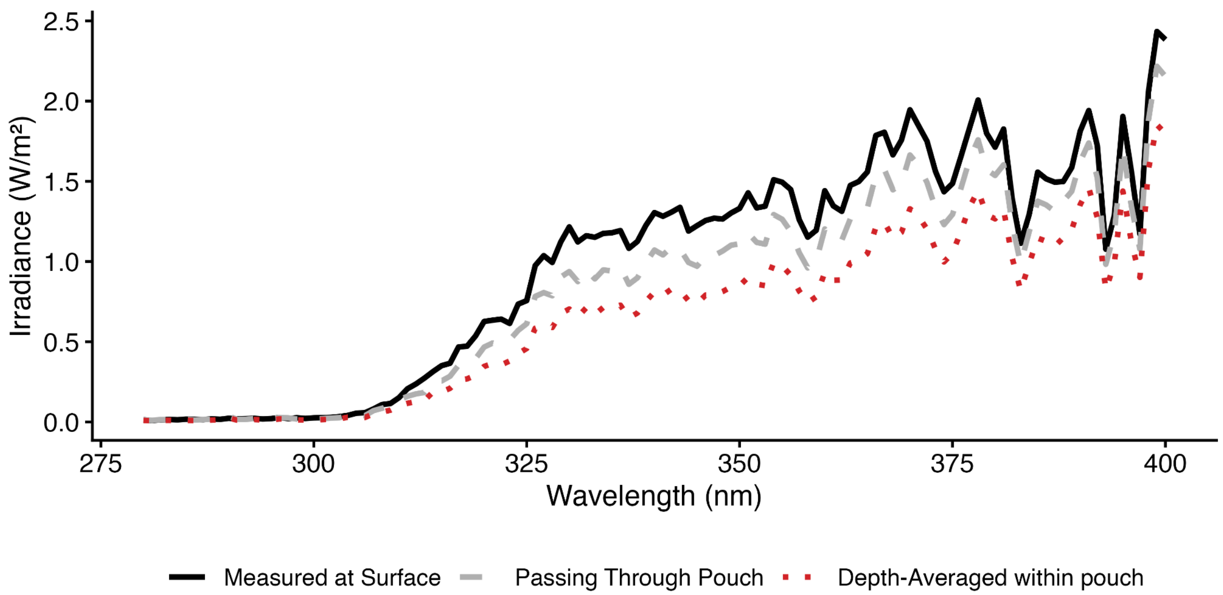

**Figure S5:** For a given radiometer measurement (15 September 2025, 0907, light condition), the measured and corrected irradiance values by wavelength

**Table S1:** Daily average UVA and UVB irradiance measured at the surface of the pouch, passing through the pouch (accounting for attenuation by the pouch material), and depth-averaged within the pouch (accounting for attenuation by the pouch material and absorption by the experimental solution).

| | | Irradiance (W/m$^2$) | | | | | |
|---|---|---|---|---|---|---|---|
| | | UVA | | | UVB | | |
| | | Measured at Surface | Passing Through Pouch | Depth-Averaged within pouch | Measured at Surface | Passing Through Pouch | Depth-Averaged within pouch |
| Day 1 | Treatment | 11.6 | 9.93 | 7.93 | 0.31 | 0.25 | 0.18 |
| | Control | 0.05 | 0.04 | 0.03 | 0.01 | 0.01 | 0.01 |
| Day 2 | Treatment | 4.0 | 3.44 | 2.75 | 0.11 | 0.09 | 0.07 |
| | Control | 0.04 | 0.04 | 0.03 | 0.01 | 0.01 | 0.01 |
| Day 3 | Treatment | 3.74 | 3.21 | 2.57 | 0.10 | 0.09 | 0.06 |
| | Control | 0.06 | 0.05 | 0.04 | 0.02 | 0.01 | 0.01 |
| Day 4 | Treatment | 2.64 | 2.27 | 1.81 | 0.07 | 0.06 | 0.04 |
| | Control | 0.04 | 0.04 | 0.03 | 0.01 | 0.01 | 0.01 |
| **Average** | **Treatment** | 5.49 | 4.71 | 3.76 | 0.15 | 0.12 | 0.09 |
| | **Control** | 0.05 | 0.04 | 0.03 | 0.01 | 0.01 | 0.01 |

### 4. Inhibition Testing

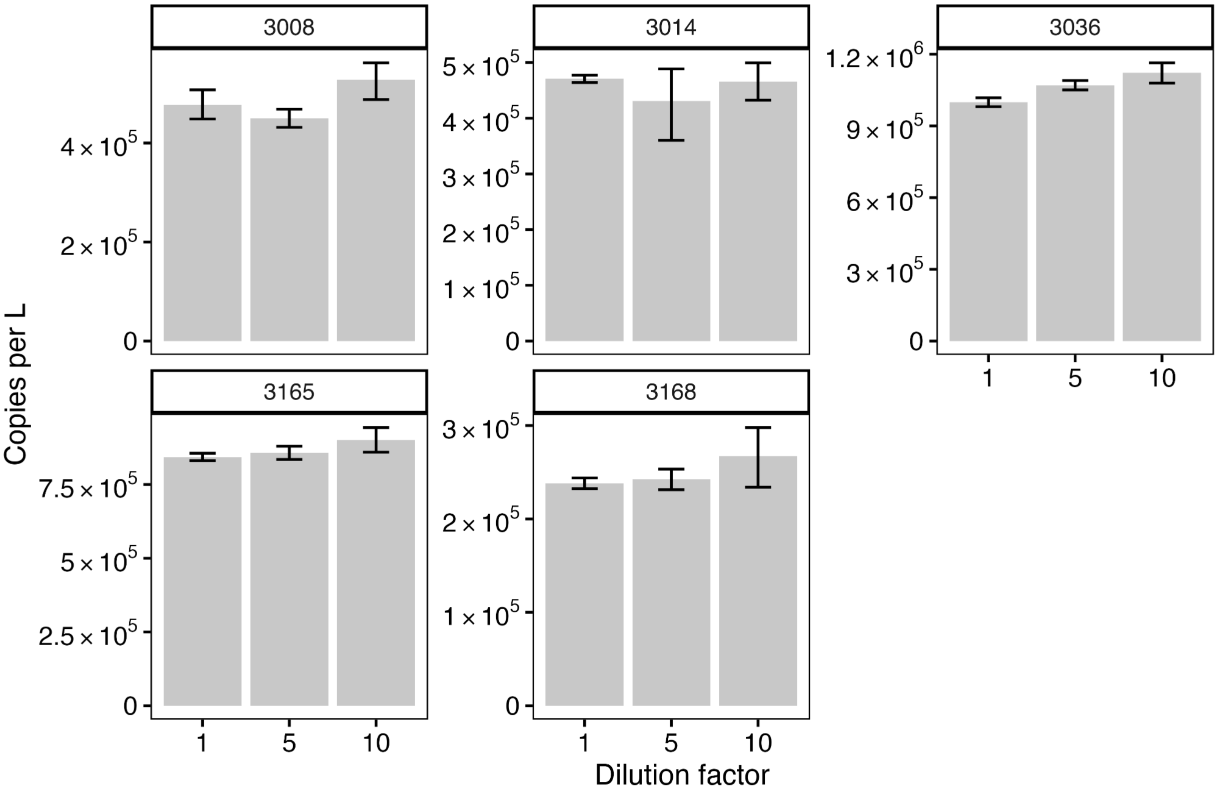

**Figure S6**: Resulting eDNA concentrations (copies/L) from inhibition testing of 5 high-concentration shedding and decay samples at 1:1, 1:5, and 1:10 dilutions (accounting for the dilution factor)

### 5. References

(1) Maraccini, P. A.; Mattioli, M. C. M.; Sassoubre, L. M.; Cao, Y.; Griffith, J. F.; Ervin, J. S.; Van De Werfhorst, L. C.; Boehm, A. B. Solar Inactivation of Enterococci and Escherichia Coli in Natural Waters: Effects of Water Absorbance and Depth. *Environ. Sci. Technol.* **2016**, *50* (10), 5068–5076. https://doi.org/10.1021/acs.est.6b00505

(2) Anderson, C. E.; Boehm, A. B. Sunlight Inactivation of Enveloped Viruses in Clear Water. *Environ. Sci. Technol.* **2023**, *57* (50), 21395–21404. https://doi.org/10.1021/acs.est.3c06680
