## Supporting Information 2 - EMMI Checklist for "Sunflower sea star (*Pycnopodia helianthoides*) environmental DNA (eDNA) shedding and decay: Influence of sunlight and comparison to other marine organisms"

| Study Description | Environmental Sampling | Sample Treatment | Sample Reduction | Nucleic-acid Extraction | Reverse Transcription | PCR Amplification | Analysis |
|---|---|---|---|---|---|---|---|
| Sunflower sea star (*Pycnopodia helianthoides*) environmental DNA (eDNA) shedding and decay: Influence of sunlight and comparison to other marine organisms<br>13 August 2026<br>Completed by Meghan M. Shea | 1 liter (decay experiment) or 0.53 liter (shedding experiment) seawater samples collected in enteral feeding pouches and gravity filtered onto 0.22 µm pore size PVDF Sterivex cartridge; described in further detail in methods | NA | NA | Extracted within 4 months of collection using a modified protocol for begining extraction inside the Sterivex cartridge (Spens et al. 2017) with the Qiagen DNeasy Blood and Tissue Kit; described in further detail in the methods | NA | Quantified using droplet digital PCR (ddPCR) and a species-specific hydrolysis probe assay for *P. helianthoides*, using 5 ul of template per reaction in duplicate reactions; described in further detail in the methods | Concentrations converted to copies of L of the original water sampled to enable calculation of decay rate constants and shedding rates; described in further detail in the methods |

### Control Checklist

| Control Checklist | Environmental Sampling | Sample Treatment | Sample Reduction | Nucleic-acid Extraction | Reverse Transcription | PCR Amplification | |
|---|---|---|---|---|---|---|---|
| Step performed | ☑ | ☐ | ☐ | ☑ | ☐ | ☑ | |
| Step has control info | ☑ | ☐ | ☐ | ☑ | ☐ | ☑ | Negative controls |
| # of control replicates | Shedding Experiment: 1 sampling point per day (4 total); Decay Experiment: 1 sampling point per day per condition (9 total) | | | 1 per extraction batch (9 total) | | 1 per ddPCR plate (9 total) | |
| Control result reported | ☑ | ☐ | ☐ | ☑ | ☐ | ☑ | |
| Method for handling failed controls described | ☐ | ☐ | ☐ | ☐ | ☐ | ☐ | |
| Added notes | No failed controls | | | No failed controls | | No failed controls | |
| Step has control info | ☐ | ☐ | ☐ | ☐ | ☐ | ☑ | Positive controls |
| Control identity described | ☐ | ☐ | ☐ | ☐ | ☐ | ☑ | |
| Control quantification method described | ☐ | ☐ | ☐ | ☐ | ☐ | ☑ | |
| # control replicates | | | | | | 1 per ddPCR plate (9 total) | |
| Control result reported | ☐ | ☐ | ☐ | ☐ | ☐ | ☑ | |
| Method for handling failed controls described | ☐ | ☐ | ☐ | ☐ | ☐ | ☐ | |
| Added notes | | | | | | No failed controls | |

### Process checklist

| Environmental Sampling | |
|---|---|
| Sample procedure | ☑ |
| Number of samples | ☑ |
| Sample amount, mean, range | ☑ |
| Sampling locations, dates, times | ☑ |
| Sample storage conditions | ☑ |

| Sample Treatment | Not applicable for this study |
|---|---|
| Treatment procedure | ☐ |
| Reagents | ☐ |

| Sample Reduction | Not applicable for this study |
|---|---|
| Reduction procedure | ☐ |
| Reagents | ☐ |
| Concentration factor | ☐ |

| Nucleic-acid Extraction | |
|---|---|
| Extraction procedure | ☑ |
| Volume or mass extracted, volume or mass obtained | ☑ |
| Extract storage conditions | ☑ |

| Reverse Transcription | Not applicable for this study |
|---|---|
| One- or two-step | ☐ |
| cDNA storage conditions (if 2 step) | ☐ |
| Reaction temperatures and times | ☐ |
| Reaction reagents and concentrations | ☐ |
| Priming method | ☐ |
| Reaction volume, added template amount | ☐ |
| RT efficiency assessment procedure (if 2-step) | ☐ |
| RT control description (if two-step) | ☐ |
| RT efficiency reported (if 2-step) | ☐ |

| qPCR or dPCR | |
|---|---|
| Target gene name, amplicon length | ☑ |
| Thermocycling temp and times | ☑ |
| Master mix composition: vendors, concentrations | ☑ |
| Additives: vendors, composition | ☑ |
| Template amount added, pre-treatment (if any) | ☑ |
| Primers: sequences, concentrations, vendors, references | ☑ |
| Amplicon confirmation method (probe, melt curve details, etc) | ☑ |
| Probe sequence, concentration, vendor, reference | ☑ |
| Instrumentation | ☑ |
| Inhibition assessment procedure | ☑ |
| Inhibition control description (if used) | ☐ |
| Number of samples tested and found inhibited | ☑ |
| Equivalent volume of sample analyzed | ☑ |

| Analysis- dPCR | |
|---|---|
| Threshold settings | ☑ |
| Technical replicates, number, well merging | ☑ |
| Partitions measured, number, mean, variance | ☑ |
| Partition volume | ☑ |
| Target copies per partition, mean, variance | ☑ |
| Program used for dPCR analysis | ☑ |
| Explanation of control results, example plots | ☑ |

| Analysis- qPCR | Not applicable for this study |
|---|---|
| Technical replicates, number, calculations | ☐ |
| Calibration standards, description, source | ☐ |
| Method of quantifying standards | ☐ |
| Calibration curve slope | ☐ |
| Calibration curve R2 | ☐ |
| Lowest standard measured or 95% LOD | ☐ |
| Cq value determination methods | ☐ |

Note to users: This checklist is provided as guidance for best practices for reporting, but is not meant to be prescriptive. Not all items in the check list will apply to all studies. Please see Borchardt et al. The Environmental Microbiology Minimum Information (EMMI) Guidelines: qPCR and dPCR Quality and Reporting for Environmental Microbiology. Environmental Science & Technology, 2021, 55, 15, 10210–10223.

Version 3.0
This version maintained by Borchardt, Boehm, Salit, Noble, Wiggington, Spencer
Date: 18 October 2023
