## Supporting Information 3 - dMIQE Checklist for "Sunflower sea star (*Pycnopodia helianthoides*) environmental DNA (eDNA) shedding and decay: Influence of sunlight and comparison to other marine organisms"

| ITEM TO CHECK | PROVIDED | COMMENT |
|---|---|---|
| | Y/N | |
| **1. SPECIMEN** | | |
| Detailed description of specimen type and numbers | Y | |
| Sampling procedure (including time to storage) | Y | Specific sampling procedure varied by experiment, but generally involved collecting water samples and gravity filtering onto Sterivex cartridges; detailed in Methods (Section 3.2, 3.3, and 3.4) |
| Sample aliquotation, storage conditions and duration | Y | Samples stored at -20°C for no more than 4 months before extraction; detailed in Methods (Section 3.4 and 3.5) |
| **2. NUCLEIC ACID EXTRACTION** | | |
| Description of extraction method including amount of sample processed | Y | Modified DNeasy Blood & Tissue Kit protocol; detailed in Methods (Section 3.5) and cited protocol |
| Volume of solvent used to elute/resuspend extract | Y | 100 µL of Buffer AE; detailed in Methods (Section 3.5) and cited protocol |
| Number of extraction replicates | Y | No extracton replication; detailed in Methods (Section 3.5) and cited protocol |
| Extraction blanks included? | Y | 1 extraction blank per extraction batch; detailed in Methods (Section 3.5) |
| **3. NUCLEIC ACID ASSESSMENT AND STORAGE** | | |
| Method to evaluate quality of nucleic acids | N | Quality was not assessed before ddPCR quantification. |
| Method to evaluate quantity of nucleic acids (including molecular weight and calculations when using mass) | N | Quantity was not assessed before ddPCR quantification. |
| Storage conditions: temperature, concentration, duration, buffer, aliquots | Y | All extracts were kept on ice until used as template in PCR, typically within 2 hours (PCR was run directly after DNA extraction); detailed in Methods (Section 3.5) |
| Clear description of dilution steps used to prepare working DNA solution | Y | Template was used undiluted except for inhibition testing; detailed in Methods (Section 3.5) |
| **4. NUCLEIC ACID MODIFICATION** | NA | |
| Template modification (digestion, sonication, pre-amplification, bisulphite etc.) | | |
| Details of repurification following modification if performed | | |
| **5. REVERSE TRANSCRIPTION** | NA | |
| cDNA priming method and concentration | | |
| One or two step protocol (include reaction details for two step) | | |
| Amount of RNA added per reaction | | |
| Detailed reaction components and conditions | | |
| Estimated copies measured with and without addition of RT* | | |
| Manufacturer of reagents used with catalogue and lot numbers | | |
| Storage of cDNA: temperature, concentration, duration, buffer and aliquots | | |
| **6. dPCR OLIGONUCLEOTIDES DESIGN AND TARGET INFORMATION** | | |
| Sequence accession number or official gene symbol | N | We are using a previously developed assay in this manuscript; this is elaborated in separate cited assay development manuscript. |
| Method (software) used for design and *in silico* verification | N | We are using a previously developed assay in this manuscript; this is elaborated in separate cited assay development manuscript. |
| Location of amplicon | Y | NADH dehydrogenase 5 (nad5) gene; detailed in Methods (Section 3.5) |
| Amplicon length | Y | 123 base pair fragment; detailed in Methods (Section 3.5) |
| Primer and probe sequences (or amplicon context sequence)** | Y | Forward primer: Pycno-nad5-v1-F (CTAACATCATTAACTTTCAGAACACG), Reverse primer: Pycno-nad5-v1-R (GCCTGTTTGGGTTAGCTGG), Probe: Pycno-nad5-v1-probe (CCAGGGATGAAATGAAACTGTAGGTGCCC); detailed in Methods (Table 1) |
| Location and identity of any modifications | N | Not applicable; no modifications |
| Manufacturer of oligonucleotides | Y | IDT; detailed in Methods (Section 3.5) |
| **7. dPCR PROTOCOL** | | |
| Manufacturer of dPCR instrument and instrument model | Y | QX200 Droplet Digital PCR System (Bio-Rad, Hercules, CA, USA); detailed in Methods (Section 3.5) |
| Buffer/kit manufacturer with catalogue and lot number | Y | Bio-Rad ddPCR Supermix for probes, no dUTP (catalog number 186-3024, Bio-Rad, Hercules, CA, USA) came from lots 64705079 and 64707759; detailed in |
| Primer and probe concentration | Y | 900 nM forward and reverse primers and 250 nM probe; detailed in Methods (Section 3.5) |
| Pre-reaction volume and composition (incl. amount of template and if restriction enzyme added) | Y | The 20 µl reaction mixture consisted of 5 µL template DNA, 1 µL of a primer and probe pre-mix (900 nM forward and reverse primers and 250 nM probe; IDT, Coralv |
| Template treatment (initial heating or chemical denaturation) | Y | Thermal conditions for the ddPCR amplifications were an initial denaturation for 10 minutes at 95°C followed by 45 cycles of 15 seconds at 95°C and 60 seconds at 60°C; detailed in Methods (Section 3.5) |
| Polymerase identity and concentration, Mg++ and dNTP concentrations*** | N | The composition of Bio-Rad ddPCR Supermix for probes, no dUTP (catalog number 186-3024, Bio-Rad, Hercules, CA, USA) is proprietary. |
| Complete thermocycling parameters | Y | Thermal conditions for the ddPCR amplifications were an initial denaturation for 10 minutes at 95°C followed by 45 cycles of 15 seconds at 95°C and 60 seconds at 6 |
| **8. ASSAY VALIDATION** | | |
| Details of optimisation performed | N | We are using a previously developed assay in this manuscript; this is elaborated in separate cited assay development manuscript. |
| Analytical specificity (vs. related sequences) and limit of blank (LOB) | N | We are using a previously developed assay in this manuscript; this is elaborated in separate cited assay development manuscript. |
| Analytical sensitivity/LoD and how this was evaluated | Y | Limits of detection were calculated for each experiment using a positive-droplet threshold as well as analytically via multiple methods; detailed in Results (Section 4.1) and also elaborated in separate cited assay development manuscript |
| Testing for inhibitors (from biological matrix/extraction) | Y | We conducted inhibition testing on a small subset of representative high concentration samples (5), by comparing their concentrations undiluted and at 1:5 and 1:10 dilutions; we did not detect evidence of inhibition (detailed in Methods "DNA Extraction and Quantification") |
| **9. DATA ANALYSIS** | | |
| Description of dPCR experimental design | Y | |
| Comprehensive details negative and positive of controls (whether applied for QC or for estimation of error) | Y | |
| Partition classification method (thresholding) | Y | |
| Examples of positive and negative experimental results (including fluorescence plots in supplemental material) | Y | Examples of positive and negative results shown in fluoresence plot included in Supporting Information (Figure S1) |
| Description of technical replication | Y | Sample extracts were amplified in duplicate; detailed in Methods ("DNA Extraction and Quantification") |
| Repeatability (intra-experiment variation) | Y | Variation among replicates reported in Figures and Results section. |
| Reproducibility (inter-experiment/user/lab etc. variation ) | N | Not assessed in this study; all samples processed in the same laboratory. |
| Number of partitions measured (average and standard deviation ) | Y | Average and standard deviation of number of partitions included for both shedding and decay experiment; detailed in Supplemental Materials |
| Partition volume | Y | 0.00085 µL; detailed in Supplemental Materials |
| Copies per partition (λ or equivalent ) (average and standard deviation) | Y | Lambda included for both shedding and decay experiment; detailed in Supplemental Materials |
| dPCR analysis program (source, version) | Y | QX Manager (version 2.2.0; Bio-Rad, Hercules, CA, USA); detailed in Methods ("DNA Extraction and Quantification") |
| Description of normalisation method | | |
| Statistical methods used for analysis | Y | A variety of statistical methods were used; detailed in Methods ("Data Analysis") and fully reproducible using published code |
| Data transparency | raw data uploaded to online repository with ID: | |

**Table S1.** dMIQE2020 checklist for authors, reviewers and editors. Authors should fill detail whether information is provided. Where 'yes' is selected use comment box to detail location of information or to include the information. Where 'no' is selected use comment box to outline rationale for omission. Sections 4 and 5 may not apply depending on experiment.

* Assessing the absence of DNA using a no RT assay (or where RT has been inactivated) is essential when first extracting RNA. Once the sample has been validated as DNA-free, inclusion of a no-RT control is desirable, but no longer essential.

** Disclosure of the primer and probe sequence is highly desirable and strongly encouraged. However, since not all commercial pre-designed assay vendors provide this information when it is not available assay context sequences must be submitted (Bustin et al. Primer sequence disclosure: A clarification of the miqe guidelines. Clin Chem 2011; 57:919-21.)

*** Details of reaction components is highly desirable, however not always possible for commercial disclosure reasons. Inclusion of catalogue number is essential where component reagent details are not available.
